# Variation at the R181 residue of p53 confers loss of p53 DNA binding cooperativity with the retention of mitochondrial-associated apoptosis

**DOI:** 10.1101/2025.08.12.669925

**Authors:** Renyta Moses, Alexandra Indeglia, Alison S Levine, Ryan Hausler, Gregory Kelly, Sven A Miller, Isabel Anez, Melissa Heller, Rosella Delgado, Caitlin Orr, Wendy Kohlmann, Anne Naumer, Jennie Vagher, Sophie R Cahill, Luke D Maese, John Karanicolas, Judy E Garber, Maureen E Murphy, Kara N Maxwell

**Affiliations:** Department of Medicine, Division of Hematology-Oncology, Perelman School of Medicine, University of Pennsylvania, Philadelphia, Pennsylvania; Graduate Group in Cell and Molecular Biology, Perelman School of Medicine, University of Pennsylvania, Philadelphia, Pennsylvania; Graduate Group in Biochemistry and Molecular Biophysics, Perelman School of Medicine, University of Pennsylvania, Philadelphia, Pennsylvania; Program in Molecular and Cellular Oncogenesis, Wistar Institute, Philadelphia, Pennsylvania; Dana-Farber Cancer Institute, Harvard Medical School, Boston, Massachusetts; Cancer Signaling and Microenvironment Program, Fox Chase Cancer Center, Philadelphia, Pennsylvania; Huntsman Cancer Institute, University of Utah, Salt Lake City, Utah; Abramson Cancer Center, Perelman School of Medicine, University of Pennsylvania, Philadelphia, Pennsylvania

**Keywords:** TP53, p53 tumor suppressor, Familial and hereditary cancers, Li Fraumeni Syndrome

## Abstract

The p53 tumor suppressor binds DNA cooperatively as a tetramer, mediated by salt-bridge interactions between p53 residues E180 and R181. Variants at the R181 residue are one of the most identified *TP53* pathogenic variants by germline genetic testing. We show that families with *TP53* p.R181H and p.R181C variants have an attenuated cancer risk phenotype compared to patients with hotspot dominant negative loss of function *TP53* variants. Despite this phenotype, we find that p53 R181H and R181C variants have significantly reduced ability to bind to p53 promoter/enhancer target sequences and transactivate p53 target genes, similar to null variants. However, p53 R181H and R181C retain wild-type p53 structure and tetramerization. In addition, R181-mutant cells undergo apoptosis through wild-type p53 activity at the mitochondria. These results indicate that retention of transcription-independent p53 tumor suppressor function results in a reduced penetrance cancer risk syndrome in humans.

**Statement of Significance:** Inherited *TP53* variants cause Li-Fraumeni Syndrome (LFS), which has significant phenotypic heterogeneity in lifetime cancer risk. Our study shows that retention of transcription-independent p53 functions, despite loss of p53 transactivation activity, results in a reduced penetrance phenotype. Classification of *TP53* variants should incorporate assays of transcriptional and non-transcriptional functions.

## Introduction

Approximately half of all human cancers acquire a mutation in *TP53*, which encodes the tumor suppressor p53(1,2). p53 is a transcription factor that transactivates genes involved in cell cycle arrest, senescence, apoptosis, ferroptosis, and other tumor-suppressive pathways(3). Germline-inactivating mutations in *TP53* occur in individuals with Li-Fraumeni Syndrome (LFS)(4), an autosomal dominantly inherited cancer predisposition condition where patients develop myriad tumors, most commonly of the brain, breast, bone, and adrenal cortex. Lifetime cancer risk in LFS patients is approximately 70-75% for males and 90-95% for females(5,6) with over half of LFS-driven tumors presenting before the fourth decade. There is high phenotypic heterogeneity between *TP53* variants leading to a large range of penetrance that is not well understood(7).

There are currently no effective precision medicine approaches for prevention or treatment of either inherited or acquired p53 mutant cancers.

The majority of both acquired and inherited mutations in p53 are missense mutations that impair p53 function and lead to increased levels of mutant protein in the cell. There are several well-established classes of cancer-associated p53 missense mutations. Structural mutations such as R175H reduce p53 stability and promote misfolding of the protein(8). Locally misfolded p53 mutations like Y220C(8) create small pockets of misfolding, which impairs p53 function. DNA contact mutants like R273H tend to retain structural folding but interfere with DNA binding(9,10). Finally, oligomerization mutants such as R337H fail to form p53 tetramers(11). A unique class of p53 mutations occurring at residues glutamic acid (E)180 and arginine (R)181 affect cooperative binding of p53 to DNA(12). Structural studies show these residues do not directly contact DNA or contribute to the structural integrity of the DNA binding domain of p53, but instead prevent p53 tetramers from cooperatively binding to DNA by disrupting intra-tetramer salt bridge formation(13,14). Engineered charge-inverting cooperativity mutations such as R181E and E180R, which have not been reported in human tumors, have been found to impair the ability of p53 to transactivate genes involved in apoptosis but retain ability to transactivate cell cycle arrest genes(15–17).

Both inherited and acquired p53 variants are thought to lead to cancer formation predominantly due to loss of the mutant ability to perform transcriptional activation; however p53 has transcription-independent tumor suppressive functions as well(18–21). A recent study of one family with the A347D mutant suggested a separation of transcriptional dependent and independent p53 functions by some hypomorphic mutants(22). Herein, we characterize naturally occurring p53 cooperativity mutants R181H and R181C that are presently reported as variants with conflicting classifications of pathogenicity and are commonly found in germline genetic testing(23). We show that these mutations have reduced DNA binding and transactivation of all p53 target genes while maintaining transcription-independent apoptotic functions. By better understanding this class of mutants, we can gain insight into key p53 functions in tumor suppression and show the importance of considering all p53 functions when inferring *TP53* variants’ potential impact on cancer risk.

## Results

### R181 variants are common and predispose individuals to numerous cancers

We reviewed 415 LFS families from research databases at three academic centers. Variation at the R181 residue (*TP53* c.542G>A;p.R181H and *TP53* c.541C>T;p.R181C) were the most common germline *TP53* variants identified in this cohort (36 of 415, 8.7% of families) (**Fig S1A-C**). After exclusion of carriers of other high risk mutations, we analyzed the cancer spectrum among 133 R181-variant carriers in 34 families (112 R181H in 27 families and 21 R181C in 7 families) (**Fig. S1D, Table S1**). Overall, 69 (51.8%) individuals had a cancer history with 109 tumors overall (**Fig. S1E)**. Breast cancer was the most frequently observed tumor type accounting for 40.4% of the total tumors, followed by skin cancer (nine cases of cutaneous melanoma and one case of Merkel skin cancer) (9.2%), thyroid cancer (8.3%), prostate cancer (7.3%), hematologic malignancies (5.5%), lung cancer (5.5%), and brain tumors (4.6%).In contrast, a different distribution of tumor types was seen in R181-mutated tumors in the AACR GENIE cohort(24), which does not differentiate between germline and somatic *TP53* mutations; the vast majority of GENIE *TP53* findings are likely to be somatic. The most common tumor types in AACR GENIE with R181 mutations were lung and colorectal cancers (**Fig. S1F**).

### R181 variants have a hypomorphic clinical phenotype

We next compared the tumor distribution in R181 carriers to tumors from patients with dominant negative/loss of function (DN/LOF) and hypomorphic variants in our cohorts (**Fig. S1G-I**).

Among R181 carriers, 42% of females had a history of breast cancer, compared to 59% of DN/LOF carriers (p=0.0085) and 38% of hypomorphic variant carriers (p=0.5515) (**Fig. 1A).** Other LFS tumors such as soft-tissue sarcoma, bone cancer, and brain tumors were less frequent in R181 variant carriers compared to DN/LOF and found at similar rates to other hypomorphic variant carriers (**Fig. 1A**). Adrenal cortical cancers and leukemias were found at similar rates between mutation types (**Fig. 1A**). The age of onset of all primary tumors in R181H and R181C carriers was significantly older compared to DN/LOF carriers (median 47 vs 30, p<0.0001) and similar to that of hypomorphic variant carriers (median 47 vs 37, p=0.0424) (**Fig 1B**, **Table S2**). The age of onset of breast cancer in R181H and R181C female carriers was also significantly older compared to DN/LOF carriers (median 43 vs 32, p<0.0001) but similar to hypomorphic variant carriers (median 43 vs 43, p=0.1375) (**Fig 1C**, **Table S2**). Compared to data from the National Cancer Institute Surveillance, Epidemiology, and End Results (NCI SEER) program (2018–2022)(25), the proportions of R181 carriers diagnosed with all cancer or breast cancer was shifted to younger age groups compared to the general population, similar to other hypomorphs but older than individuals with DN/LOF *TP53* variants (**Fig. 1D-E, Table S2**). Based on cancer diagnosis, age of onset, family history, phenotype characteristics, LFS tumor spectrum classifications(26) were assigned to all R181-variant carriers compared to individuals with DN/LOF and hypomorphic variants. A higher fraction of R181 carriers presented as attenuated LFS compared to DN/LOF variant (63% vs 16%, p<0.0001) and hypomorphic variant families (63% vs 36%, p<0.0001) (**Fig. 1F-G**, **Fig. S2, Table S3**).

**Figure 1.**
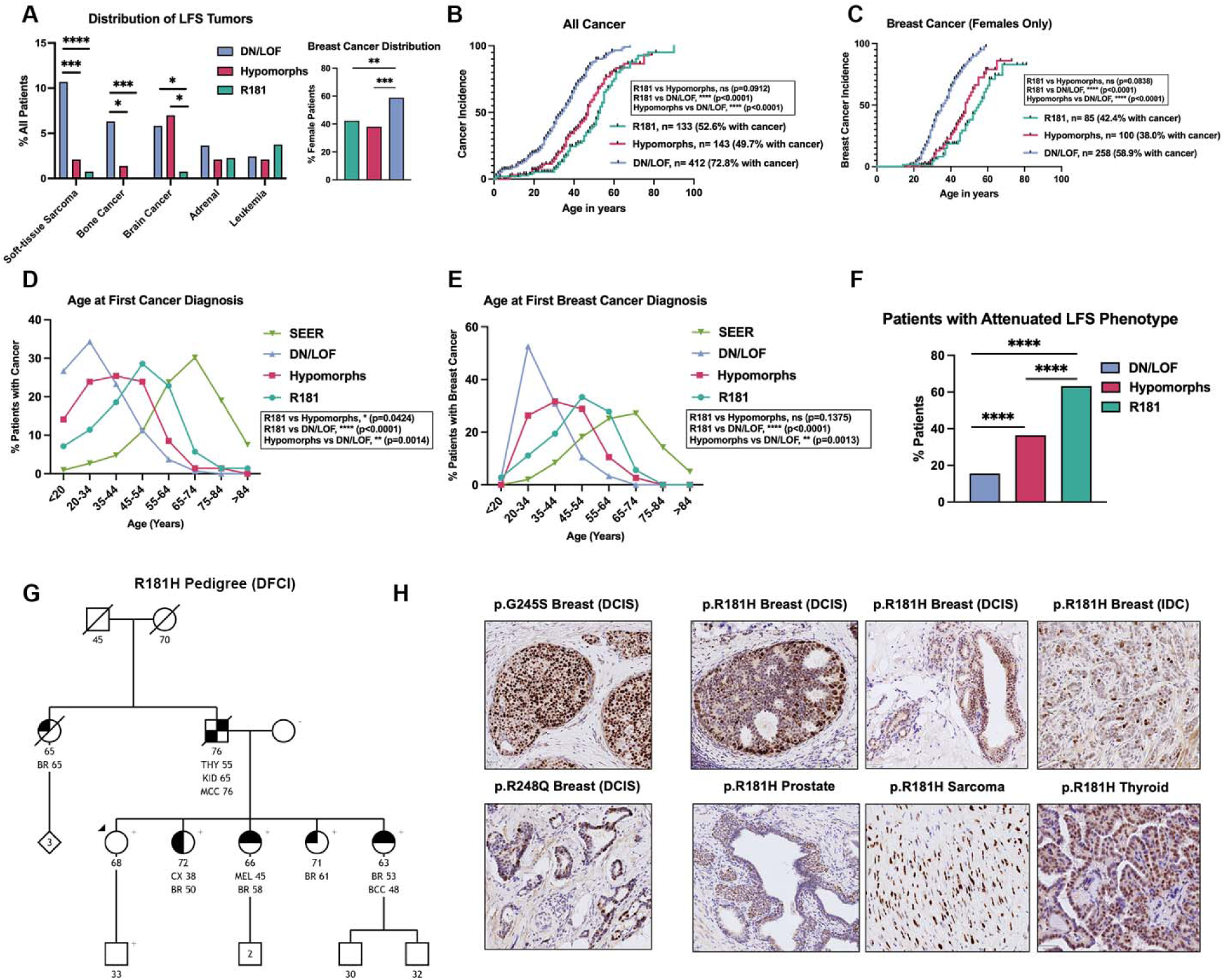
Individuals with germline R181 variants have an attenuated LFS clinical phenotype. (A) Fraction of individuals with *TP53* c.542G>A;p.R181H and *TP53* c.541C>T;p.R181C, Dominant Negative/Loss of Function (DN/LOF) variants in *TP53* or hypomorphic variants in *TP53* who had a diagnosis of breast cancer and other common LFS cancers. Variants included as hypomorphic were: p.G105D, p.R110C, p.R110L, p.T125M, p.N131K, p.G154D, p.R156H, p.R158C, p.A161T, p.R175C, p.H178Q, p.P190R, p.H193Q, p.H214Y, p.V218M, p.R267Q, p.R267W, p.C277R, p.D281E, p.R282Q, p.G245C, p.G334R, p.A347D. (B) Fraction of individuals with R181, DN/LOF or hypomorphic variants with any first cancer diagnosed from birth to age 100. (C) Fraction of female individuals with R181, DN/LOF or hypomorphic variants with a first breast cancer diagnosed from birth to age 100. (D) Of individuals with any cancer, fraction that developed any cancer by age grouping in R181, DN/LOF or hypomorphic variants compared to general population data from the National Cancer Institute Surveillance, Epidemiology, and End Results (NCI SEER) program (2018–2022). (E) Of female individuals with breast cancer, fraction that developed breast cancer by age grouping in R181, DN/LOF or hypomorphic variants compared to general population data from the National Cancer Institute Surveillance, Epidemiology, and End Results (NCI SEER) program (2018–2022). (F) Fraction of n=133 R181-variant carriers designated as attenuated LFS by revised Kratz criteria compared to fraction of n=143 carriers of hypomorphic and n=412 carriers of DN/LOF variants. (G) Representative pedigree from a family with the R181H variant. (H) Immunohistochemistry (IHC) of p53 on sections of formalin-fixed paraffin-embedded (FFPE) tumor blocks from patients with germline *TP53* variants p.G245S (breast cancer), p.R248Q (breast cancer), and p.R181H (two breast cancers, a sarcoma and a thyroid cancer). Fischer’s exact test; * P < 0.05, *** P < 0.001, **** P < 0.0001.

### R181-mutant tumors undergo *TP53* loss of heterozygosity

One of the first tumor initiating steps in LFS cancers is loss of heterozygosity (LOH) in which the WT copy of *TP53* is lost(27,28). Cells that undergo *TP53* LOH lose the functional WT p53 which is necessary to induce the expression of the negative regulator of p53, MDM2. As such, cells that become fully mutant with a missense *TP53* mutation stabilize the mutant p53 protein and can be detected through p53 immunohistochemistry (IHC), as shown for two hotspot DN/LOF mutations, G245S and a R248Q (**Fig. 1H**). Similarly, we observed positive p53 staining in six R181H tumors (n=3 breast, n=1 each thyroid, sarcoma, prostate) strongly suggesting LOH events and that R181H is the driving mutation in these tumors. Paired tumor-normal DNA sequencing from breast tumors with p.R181H (n= 2) and *TP53* loss of function variants p.G245S (n= 1), p.R248Q (n= 1) confirmed LOH at the *TP53* region on the chromosome (17p13.1) (**Fig. S3**).

### R181 variants are cooperativity mutants

Whereas tetramerization of p53 is mediated by residues in the oligomerization domain (aa. 325-355), stabilizing interactions in the DNA-binding domain (DBD) between the positively charged R181 of one p53 monomer with the negatively charged E180 of the opposite p53 monomer mediate cooperative binding of p53 to a consensus element(14) (**Fig. 2A**). AlphaFold models of p53 binding DNA suggest that R181H and R181C are unable to form the necessary salt-bridges for cooperativity (**Fig. 2B-C**). To experimentally analyze p53 cooperative binding to specific DNA sites, the purified DBD (aa 94-292) of wild-type p53 (WT), R181H, and R181C were subjected to fluorescence polarization assays. Whereas the WT p53-DBD demonstrated cooperative binding to the *CDKN1A*(p21), *BAX*, and *BBC3*(Puma) promoters (**Fig. 2D**), cooperative binding to these promoter sequences was mildly reduced in R181H (**Fig. 2E**) and more significantly reduced in R181C (**Fig. 2F**) mutant DBDs. Directly comparing the Hill slope for the *CDKN1A*(p21) promoter (**Fig. 2G**) shows a significant reduction for the R181C compared to WT DBD (**Fig. 2H**).

**Figure 2.**
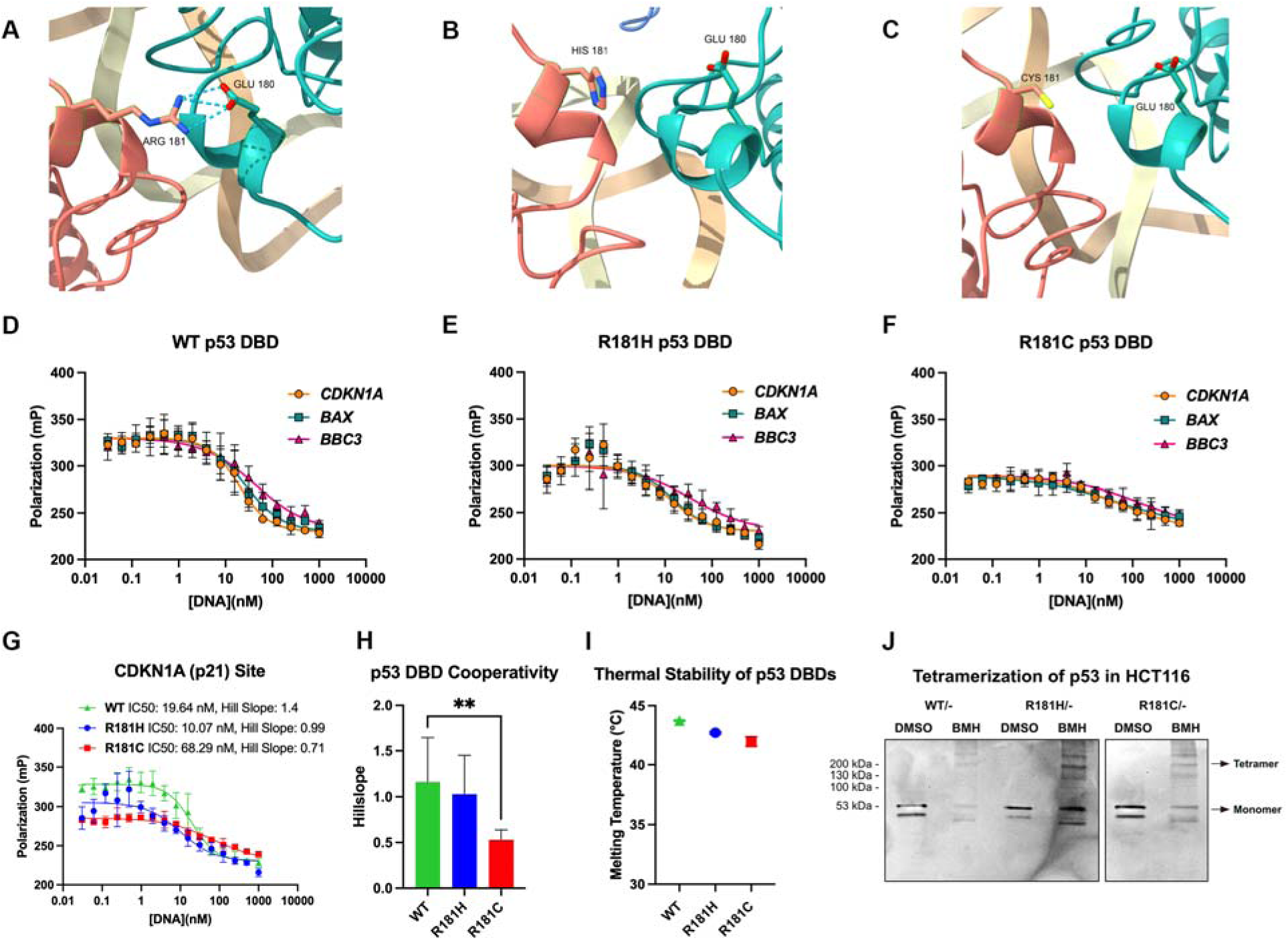
p53 R181 mutants are defective in DNA binding cooperativity. (A) AlphaFold3 model using full-length WT p53 demonstrating the E180-R181 salt-bridge between two WT p53 monomers; blue = positive charge, red = negative charge. (B) AlphaFold3 model using full-length p53-R181H demonstrating the inability of the H181 mutant monomer to form a salt-bridge with E180 residue of the other p53 monomer; blue = positive charge, red = negative charge. (C) AlphaFold3 model using full-length p53-R181C demonstrating the inability of the C181 mutant monomer to form a salt-bridge with E180 residue of the other p53 monomer; yellow = neutral charge, red = negative charge. (D) Fluorescence polarization assay of purified p53 WT DNA binding domain (DBD) subjected to p53 target promoter sites of *CDKN1A*, *BAX*, and *BBC3*(Puma). (E) Fluorescence polarization assay of purified p53 R181H DNA binding domain (DBD) subjected to p53 target promoter sites of *CDKN1A*(p21), *BAX*, and *BBC3*(Puma). (F) Fluorescence polarization assay of purified p53 R181C DNA binding domain (DBD) subjected to p53 target promoter sites of *CDKN1A*(p21), *BAX*, and *BBC3*(Puma). (G) Fluorescence polarization assay of purified p53 WT, R181H, and R181C DNA binding domain (DBD) subjected to p53 target promoter site of *CDKN1A*(p21). (H) Comparison of Hill slopes for p53 WT, R181H, and R181C DNA binding domain (DBD) to the *CDKN1A*(p21) promoter site. Data depicted in D-H are the average results of three independent experiments; student’s t-test ** P < 0.001 (I) Melting temperature of purified p53 WT, R181H, and R181C DBDs as determined by differential scanning fluorimetry. Data averaged from four independent experiments. (J) Western blot for p53 in CRISPR-engineered p53 WT/-, R181H/- and R181C/-HCT116 cells after DMSO or treatment with bismaleimidohexane (BMH).

### Structural and oligomerization ability retained in R181 variants

The purified DBD of WT p53, R181H, and R181C p53 were used to investigate structural differences in R181 variants. Using Differential Scanning Fluorimetry to measure the thermal point of denaturation of proteins, we observed WT-like (43.7°C) melting temperatures (T_m_) of the DBDs of R181H (42.7°C) and R181C (42.0°C) (**Fig. 2I**), indicating modest effects of R181 mutations on structural stability. This contrasts with known structural mutants of p53 such as the hotspot variant R175H where the T_m_ of the DBD markedly reduces to 35.0°C, as reported by other groups(29). To examine the impact of R181 mutants on oligomerization, we created CRISPR knockout lines in the HCT116 background of R181H/- and R181C/-, to mimic the tumor background. As a control, we created WT/-cells using the same procedure.

Bismaleimidohexane (BMH)-mediated crosslinking of p53 in these CRISPR-engineered p53 WT/-, R181H/- and R181C/-HCT116 cells revealed that both R181H and R181C p53 mutants form tetramers in a manner indistinguishable from WT p53 (**Fig. 2J**).

### R181 variants have reduced ability to transactivate p53 target genes

The fundamental role of p53 in tumor suppression is to transactivate a variety of genes involved in processes such as apoptosis and cell cycle arrest(30). To investigate the tumor suppressive capability of R181H and R181C mutants in a broad series of cell lines, we created CRISPR-engineered cell lines containing WT p53 or hemizygous mutant R181H and R181C in three different cancer backgrounds (HCT116, MCF7, and LNCaP). WT/-, R181H/- and R181C/- cell lines in the HCT116 and MCF7 backgrounds were treated with the p53-activating agent Nutlin-3a(31) for 0, 8 and 24 hours and subjected to RNAseq. Differential expression analysis demonstrated a number of p53 target genes(32) were significantly down-regulated in R181H/- and R181C/- cells compared to WT/- cells after 8 (**Fig. S4A**) and 24 hours (**Fig. 3A**) of nutlin in the HCT116 background. As expected, nutlin significantly induced the expression of 48 and 143 of 326 known p53 target genes(32) at 8 hours (**Fig. S4B, Table S4)** and 24 hours (**Fig. 3B, Table S5**) in WT/- HCT116, respectively. In comparison, both R181H/- and R181C/- cells showed markedly impaired transactivation of the vast majority of these p53 target genes (**Fig. 3B, Fig. S4B, Table S4-5**). Ingenuity Pathway Analysis (IPA) showed p53 pathways were among the most significantly downregulated pathways at 8 hours of nutlin treatment in R181H/- and R181C/- HCT116 cells compared to WT/- (**Fig. 3C, Table S6-7**). Regulator analysis in IPA confirmed reduced activity of *TP53* and Downstream Regulators of *TP53* (*CDKN1A*) in the R181-mutant cells (**Fig. S4C**). RNAseq data was confirmed by qPCR for canonical p53 targets *CDKN1A* (p21), *BBC3* (Puma) and *GDF15* in HCT116 cells (**Fig. S4D-F**). The reduced transactivation of p53 targets by R181H and R181C mutants was recapitulated in the MCF7 cell line (**Fig. 3D-F, Fig. S4G-I, Table S8-11**). Western blotting of canonical p53 targets MDM2 and p21 showed reduced protein induction in the R181-mutant cells in the background of HCT116, MCF7, and LNCaP (**Fig 3G-I**). In patient-derived lymphoblastoid cell lines (LCLs) that are heterozygous (WT/R181H), we observed significant levels of p53 target induction upon Nutlin-3a treatment, suggesting that the R181 mutant may lack dominant negative activity, at least in this cell type (**Fig. S4J**).

**Figure 3.**
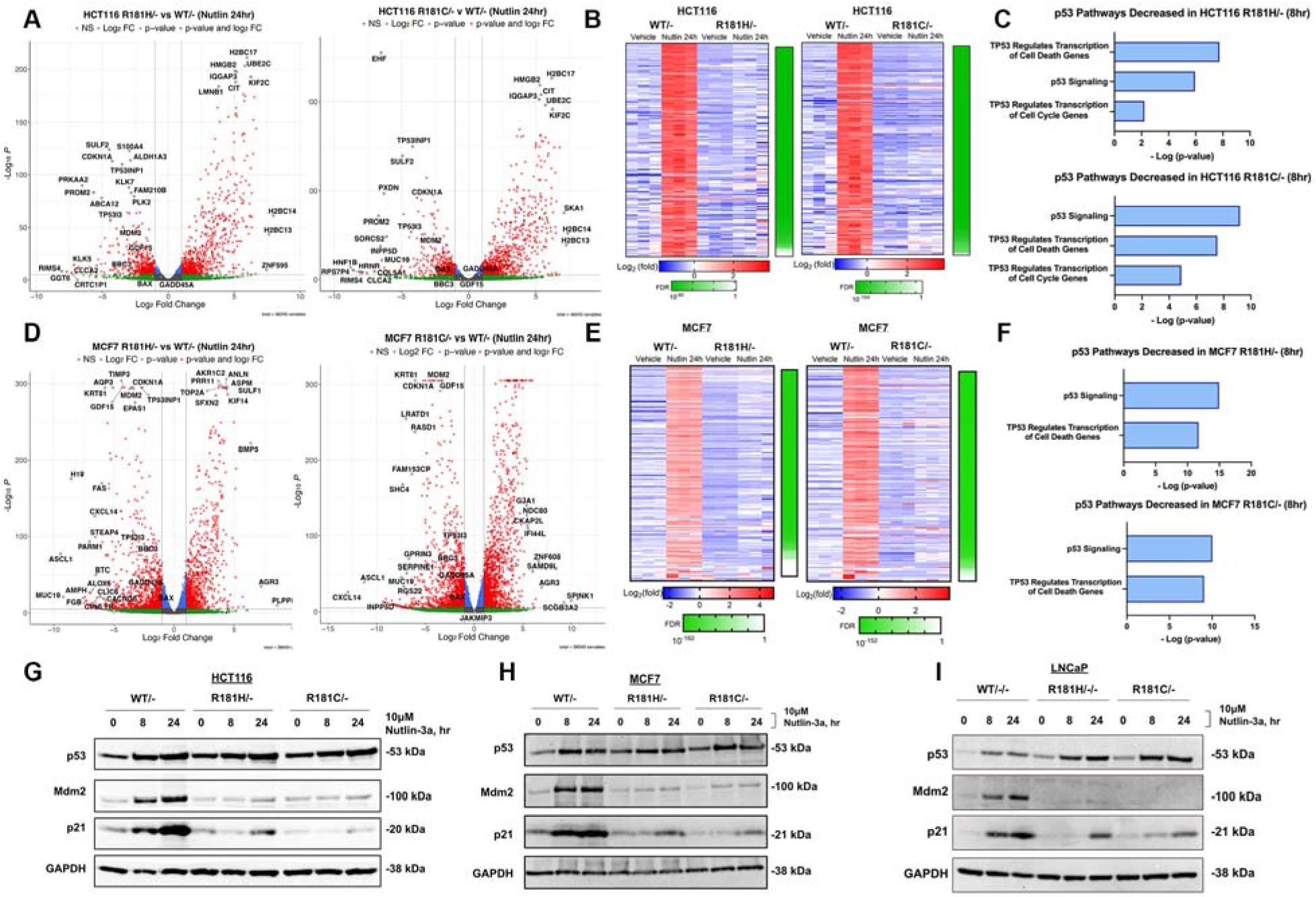
R181 variants have reduced ability to transactivate p53 target genes. (A) Volcano plots of differentially expressed genes in HCT116 R181H/- cells compared to p53 WT/- cells and HCT116 R181C/- cells compared to p53 WT/- cells after 24 hours of Nutlin-3a treatment. (B) Heatmap showing log2(Fold Change) after 24 hours of vehicle or Nutlin-3a in HCT116 p53 WT/- cells and R181H/- cells; genes shown are known p53 targets that were significantly up-regulated in WT/- cells (n=143). Bar to the right of the heatmap depicts the false discovery rate comparing the fold change between treatment and vehicle in HCT116 p53 WT/- versus R181H/- cells. Heatmap showing log2(Fold Change) after 24 hours of vehicle or Nutlin-3a in HCT116 p53 WT/- cells and R181C/- cells; genes shown are known p53 targets that were significantly up-regulated in WT/- cells (n=143). Bar to the right of the heatmap depicts the false discovery rate comparing the fold change between treatment and vehicle in HCT116 p53 WT/- versus R181H cells. (C) Bar plot showing the -log(p-value) of the most significantly decreased pathways in HCT116 R181H/- vs WT/- 8 hours after Nutlin-3a treatment. Bar plot showing the - log(p-value) of the most significantly decreased pathways in HCT116 R181C/- vs WT/- 8 hours after Nutlin-3a treatment. (D) Volcano plots of differentially expressed genes in MCF7 R181H/- cells compared to p53 WT/- cells and MCF7 R181C/- cells compared to p53 WT/- cells after 24 hours of Nutlin-3a treatment. (E) Heatmap showing log2(Fold Change) after 24 hours of vehicle or Nutlin-3a in MCF7 p53 WT/- cells and R181H/- cells; genes shown are known p53 targets that were significantly up-regulated in WT/- cells (n=152). Bar to the right of the heatmap depicts the false discovery rate comparing the fold change between treatment and vehicle in MCF7 p53 WT/- versus R181H/- cells. Heatmap showing log2(Fold Change) after 24 hours of vehicle or Nutlin-3a in MCF7 p53 WT/- cells and R181C/- cells; genes shown are known p53 targets that were significantly up-regulated in WT/- cells (n=152). Bar to the right of the heatmap depicts the false discovery rate comparing the fold change between treatment and vehicle in MCF7 p53 WT/- versus R181H/- cells. (F) Bar plot showing the -log(p-value) of the most significantly decreased pathways in MCF7 R181H/- vs WT/- 8 hours after Nutlin-3a treatment. Bar plot showing the -log(p-value) of the most significantly decreased pathways in MCF7 R181C/- vs WT/- 8 hours after Nutlin-3a treatment. (G) Western blot of p53, Mdm2, p21, and GAPDH (loading control) in HCT116 p53 WT/-, R181H/-, and R181C/- cells after 0, 8, and 24 hours of Nutlin-3a treatment. (H) Western blot of p53, Mdm2, p21, and GAPDH (loading control) in MCF7 p53 WT/-, R181H/-, and R181C/- cells after 0, 8, and 24 hours of Nutlin-3a treatment. (I) Western blot of p53, Mdm2, p21, and GAPDH (loading control) in LNCaP p53 WT/-, R181H/-, and R181C/- cells after 0, 8, and 24 hours of Nutlin-3a treatment.

### R181 variants are proliferative and unable to cell cycle arrest

To investigate the functional phenotypes associated with the gene expression changes in R181 mutant cell lines, we next performed Ki67 staining. This analysis revealed that R181 mutant cells did not decrease proliferation after p53 activation with nutlin as p53 WT/- cells did, similar to that of p53 null cells (**Fig. 4A-B**). Since R181H/- and R181C/- cells were unable to induce the p53 target cell cycle protein p21, we performed cell cycle arrest assays using propidium iodide staining of DNA; these analyses revealed a decreased ability of mutant cells to G1 cell cycle arrest upon p53 activation with nutlin as was seen for WT/- cells (**Fig. 4C**). This led to an increased percentage of R181-mutant cells in the proliferative S phase and G2/M phase after 24 hours of Nutlin-3a treatment, and decreased proportion of cells in the Sub-G1 phase after 48 hours of Nutlin-3a (**Fig. 4D-F**).

**Figure 4.**
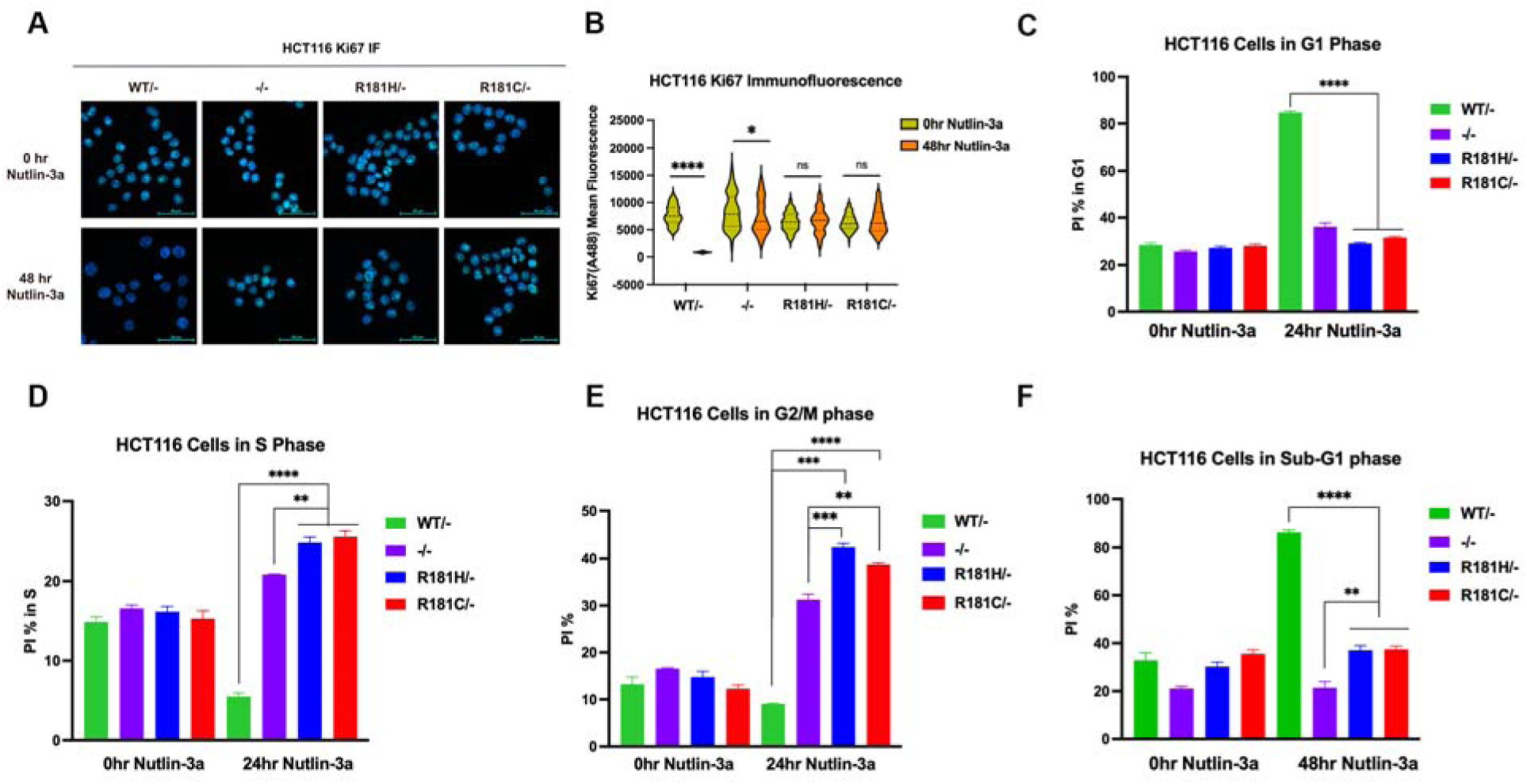
R181 variants are proliferative and unable to cell cycle arrest. (A) Representative microscopy images of Ki67 immunofluorescence (IF) of HCT116 p53 WT/-, -/-, R181H/-, and R181C/- cells at 0 and 48 hours of Nutlin-3a treatment; blue = DAPI, green = A488 (Ki67). (B) Violin plot of mean fluorescence intensity of green A488 signal (staining Ki67) of Ki67 IF in HCT116 p53 WT/-, -/-, R181H/-, and R181C/- cells at 0 and 48 hours of Nutlin-3a treatment. (C) Bar plot showing propidium iodide (PI) flow staining in G1 phase fraction (2N) in HCT116 p53 WT/-, -/-, R181H/-, and R181C/- cells at 0 and 24 hours of Nutlin-3a treatment. (D) Bar plot showing PI flow staining in S phase fraction (2N-4N) in HCT116 p53 WT/-, -/-, R181H/-, and R181C/- cells at 0 and 24 hours of Nutlin-3a treatment. (E) Bar plot showing PI flow staining in G2M phase fraction (4N) in HCT116 p53 WT/-, -/-, R181H/-, and R181C/- cells at 0 and 24 hours of Nutlin-3a treatment. (F) Bar plot showing PI flow staining in Sub-G1 phase fraction (<2N) in HCT116 p53 WT/-, -/-, R181H/-, and R181C/- cells at 0 and 48 hours of Nutlin-3a treatment. Data averaged from 3 independent experiments. Student’s t-test * P < 0.05, ** P < 0.01, *** P < 0.001, **** P < 0.0001.

### R181 variants demonstrate impaired ability to bind to p53 consensus binding sites

The *in-vitro* assays using purified DBDs of R181-mutant p53 suggested reduced cooperative binding to specific sites on DNA. To address R181 mutant binding ability in human cells, we employed chromatin immunoprecipitation-sequencing (ChIP-seq) to globally assess the ability of the R181 mutants to bind to p53 target sites on DNA. Nutlin significantly induced the binding of p53 to 2469 sites on DNA in p53 WT/- HCT116 cells. In contrast, p53 binding was essentially abolished in R181H/- and R181C/- cells (**Fig. 5A, Fig. S5A**). These sites include promoters and enhancers for important p53 targets such as *CDKN1A* (p21) and *BAX* (**Fig. 5B-C**). We performed HOMER Motif Analysis on sites with enriched binding by WT p53 compared to by R181-mutant p53. As expected, the sites significantly enriched in WT p53 over R181-mutant p53 were p53 motifs all containing the canonical p53 response element (**Fig. 5D**). Interestingly, we next used HOMER to assess the DNA binding sites retained in cells containing the R181 variants, and identified binding sites for ETS motifs (**Fig. S5B**).

**Figure 5.**
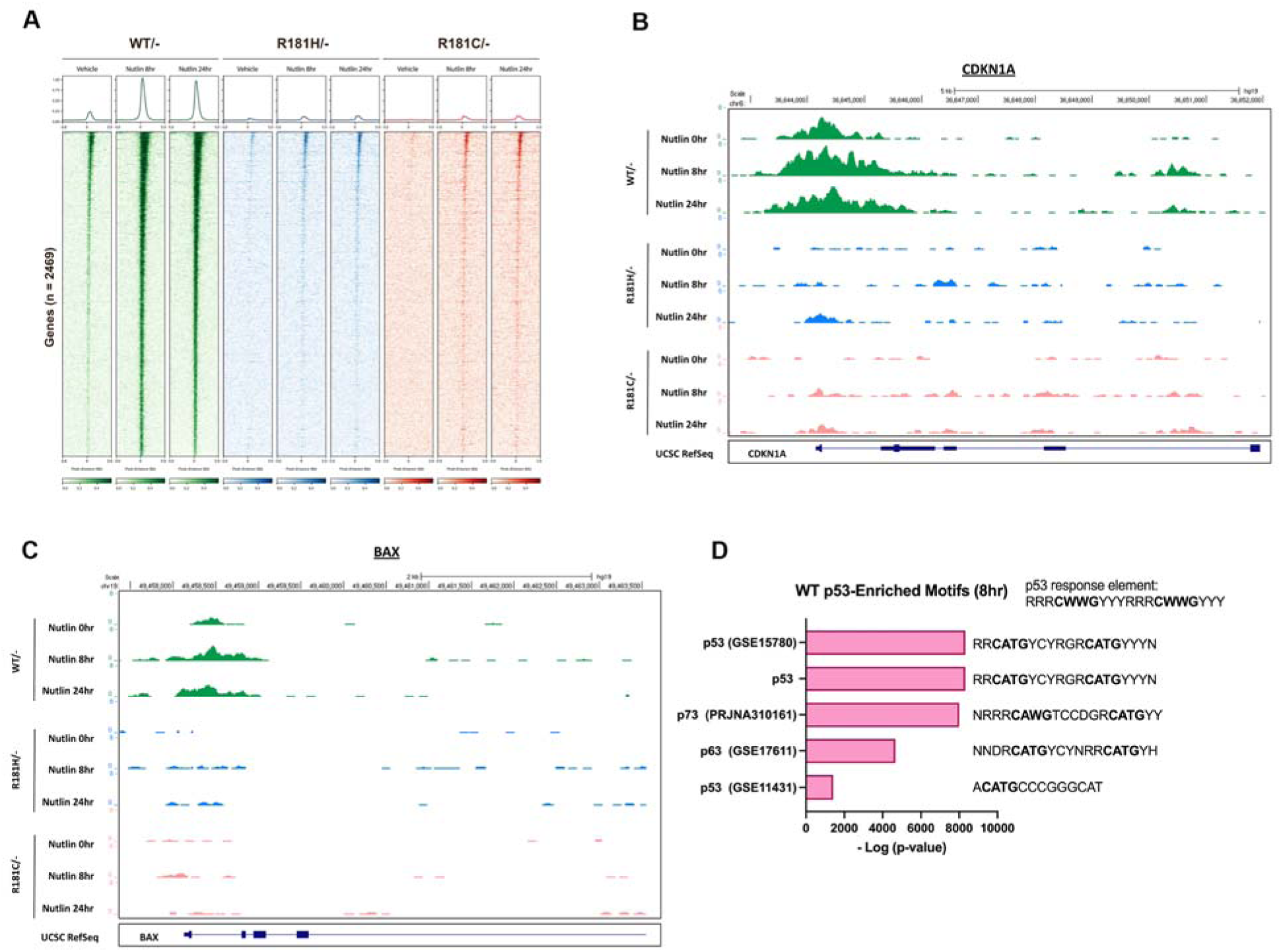
R181 variants show poor binding to p53 consensus sites. (A) Heatmap showing 2469 DNA significant peaks indicating sites bound by p53 in HCT116 p53 WT/-, R181H/-, and R181C/- cells in 0, 8, and 24 hours of Nutlin-3a treatment by chromatin immunoprecipitation sequencing (ChIP-seq) (replicate 1). (B) ChIP peaks of p53 at the *CDKN1A*(p21) target site in HCT116 p53 WT/-, R181H/-, and R181C/- cells under 0, 8, and 24 hours of Nutlin-3a treatment. (C) ChIP peaks of p53 at the *BAX* target site in HCT116 p53 WT/-, R181H/-, and R181C/- cells under 0, 8, and 24 hours of Nutlin-3a treatment. (D) Bar plot showing -log(p-value) of the most significantly enriched motifs by p53 in HCT116 WT/- cells over R181H/- and R181C/- cells. Data are representative of n = 2 independent experiments.

### R181 variants retain transcription-independent apoptosis function

Given the transcriptional defects of R181 mutants, we were surprised to find that these mutants retained the ability to suppress colony formation when transfected into H1299 (p53-null) cells, suggesting that these mutants retained some tumor suppressor function when overexpressed (**Fig. 6A**). Consistent with this, we found that subcutaneous injection of R181H/- and R181C/- HCT116 cells into the flanks of NSG SCID mice revealed similar tumor growth rates and final tumor weights compared to WT/- cells (**Fig. S6A-B**). To determine whether these variants might retain apoptotic ability, we performed Annexin V assays after treatment with 5-fluorouracil (5FU), showing comparable apoptosis levels between the R181 variants and WT p53 (**Fig. 6B**). This result was confirmed using immunoblotting of 5FU-treated cells with the apoptosis marker cleaved lamin A (CLA), which was increased in R181 cells despite an absence of induction of the transcriptional apoptotic target *BBC3* (Puma) (**Fig. 6C**). In order to determine whether the R181 mutants retained p53 transcription-independent pro-apoptotic response to DNA damage, we determined whether they could engage the direct, mitochondrial pathway of p53-mediated cell death(18). In both unstressed and 5FU treated cells, we observed no differences in mitochondrial structure measured through MitoTracker in R181 mutant or WT cells (**Fig. S7A**). Further, there were no differences in mitochondrial content measured through MitoTracker and TOMM20 flow cytometry staining, nor mitochondrial function measured through Seahorse Mito Stress Test in unstressed R181-mutant cell lines, compared to WT (**Fig. S7B-D**). We next determined whether these mutants could interact with mitochondrial BAK(19) using proximity ligation assay (PLA). This analysis revealed that both p53 R181H and R181C proteins associate with mitochondrial BAK upon DNA damage, to a level similar to WT p53 (**Fig. 6D-E**). Consistent with this finding, we found that R181-mutant cells demonstrated reduced mitochondrial cytochrome c staining upon 5FU treatment by flow cytometry, indicative of intact cytochrome c release in R181 mutant cells similar to WT (**Fig. 6F**).

**Figure 6.**
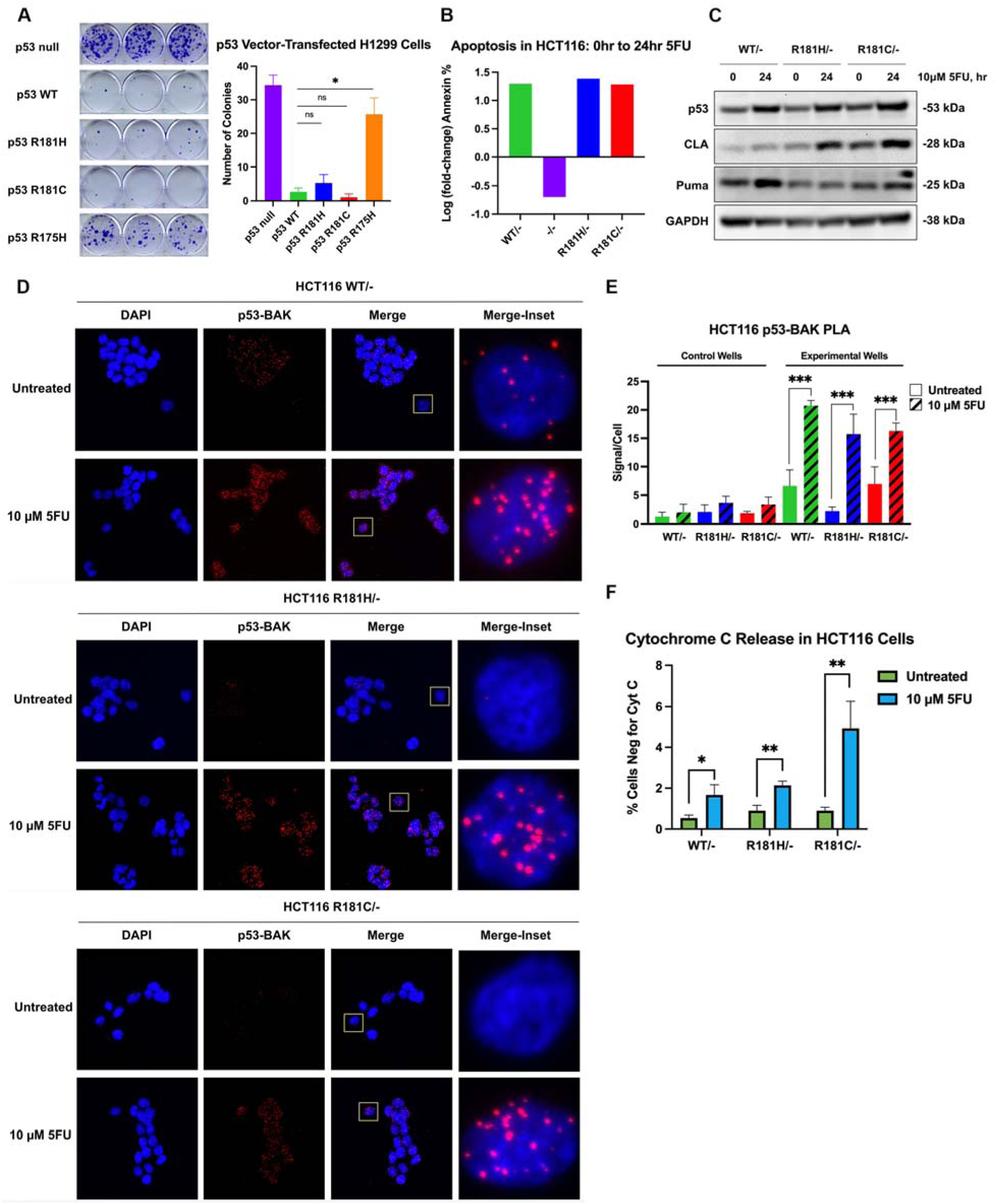
R181 variants retain p53 transcription-independent apoptosis function. (A) Images of crystal violet stained colonies of H1299 cells transfected with empty vector (p53 null), p53 WT, p53 R181H, p53 R181C, and p53 R175H CMV vectors. Bar plot quantifying colonies in H1299 cell lines transfected with empty vector (p53 null), p53 WT, p53 R181H, p53 R181C, and p53 R175H CMV vectors. (B) Bar plot showing log(fold-change) of cells staining positive for Annexin V in HCT116 p53 WT/-, R181H/-, and R181C/- from 0 to 24 hours of 5-fluorouracil (5FU) treatment. (C) Western blot of p53, cleaved lamin A (CLA), Puma, and GAPDH (loading control) in HCT116 p53 WT/-, R181H/-, and R181C/- cells in 0 and 24 hours of 5FU treatment. (D) Representative microscopy images showing proximity ligation assay (PLA) between p53 and Bak in HCT116 p53 WT/-, R181H/-, and R181C/- cells in untreated and 5FU-treated conditions; blue = DAPI, red = p53-Bak interaction. (E) Bar plot showing quantified p53-Bak PLA signals per cell in untreated and 5FU-treated HCT116 p53 WT/-, R181H/-, and R181C/- cells. (F) Bar plot showing percentage of cells negative for cytochrome c (cyt c) flow staining in HCT116 p53 WT/-, R181H/-, and R181C/- cells with and without 5FU-treatment. Data representative of three independent experiments. Student’s t-test ns P > 0.05, * P < 0.05, ** P < 0.01, *** P < 0.001.

## Discussion

Over 25,000 variants in p53 have been reported, including over 2000 inherited in humans(33). Dominant negative loss of function p53 missense mutations typically lead to classic Li Fraumeni Syndrome (LFS) with extremely high risks of cancer(34,35). However, with expanded germline genetic testing, significant phenotypic heterogeneity has been reported in individuals with germline *TP53* mutations(26). In this study, we show that the *TP53* R181H variant is the most frequently observed variant in germline carriers of *TP53* variants in three of the largest LFS research registries in the United States. Individuals with R181 variations have numerous tumors, particularly breast cancer, but fewer classic LFS tumors such as soft-tissue sarcomas, bone tumors and brain cancer, and a later age of onset of the first primary tumor.

Despite this hypomorphic clinical phenotype in R181-variant patients, our results show that p53 R181H and R181C have markedly impaired transactivation ability compared to WT p53 due to reduced p53 cooperative binding to DNA. While p53 R181 variants induced increased proliferation and were unable to induce G1 cell cycle arrest, they retained WT-like activity at the mitochondria to induce transcription-independent apoptosis. This retained tumor suppressive function, which does not rely on transactivation, may explain the attenuated LFS phenotype observed in patients.

The *TP53* R181H variant was among the variants reported in the original description of families with LFS(36). We find that R181H is the most prevalent variant in LFS families at Penn, DFCI, and HCI separately. The population in these cohorts contains significantly more individuals with R181H compared to R181C, but this may be because R181C appears to be a Palestinian founder mutation(37–40). R181H was also identified as the most frequently carried *TP53* variant in a combined population- and hospital-ascertained biobanking cohort (UK Biobank, Geisinger, and Penn)(41). At present, R181 mutants are characterized as variants with conflicting reports of pathogenicity on ClinVar(23) (R181H Accession: VCV000142320.65; R181C Accession: VCV000142624.23). Our findings reveal that individuals with these variants lack the strong cancer phenotype observed in complete loss of function variants like R175H, whose families develop many more tumors and at younger ages, and are more similar to other reported hypomorphic variants. LFS classification criteria were recently expanded to include a category of attenuated LFS(26), which we show is the predominant type of LFS observed in R181-variant carriers. Despite this attenuated phenotype, cells carrying these variants undergo LOH as an initial step in germline *TP53* variant-driven tumorigenesis. This observation strongly suggests that R181 variants are pathogenic, given that the tumors lose the WT copy and are driven by the inability of the mutant to perform specific p53 functions.

Many hypomorphic variants of p53 have been reported to retain some, if not most, levels of transcriptional activity(42); therefore, our findings of completely ablated transcriptional ability and p53 binding site activity in hemizygous R181H and R181C mutant human cancer cell lines were unexpected given the attenuated clinical phenotype and retained ability of these mutants to suppress colony formation when overexpressed. Further, our results contrast with reports on an engineered p53 mutant, E180R, which retains the ability to transactivate cell cycle arrest targets such as CDKN1A (p21) and bind the *CDKN1A* promoter but loses the ability to transactivate apoptosis targets(15–17,43). These different results could be due to prior studies using overexpression in cell lines as compared to endogenous mutagenesis, in support of this premise, recent work has demonstrated the importance of p53 dosage and the resultant amount of mutant p53 in a system, when studying p53 mutant functional effects(44).

Given that the R181 variants did not retain any transcriptional activity, we sought transcription-independent functions of p53 that could explain the attenuated clinical phenotype. p53 has been shown to induce apoptosis by localizing to the mitochondria where it activates proapoptotic BAK which leads to cytochrome c release, caspase cleavage, and cell death(18,19). This mechanism of p53-mediated apoptosis does not rely on transcriptional activity; indeed this mechanism is also retained in the LFS-associated A347D variant and the artificial mutant R181E(20,22,43). The majority of pathogenic p53 mutations are in the DNA-binding domain, highlighting the importance of p53 binding to DNA as a key transcription factor. However, our results strongly suggest that determining whether p53 mutants retain transcription-independent functions such as mitochondrial apoptosis are invaluable in variant classification and hence clinical management of patients with hypomorphic variants as compared to complete LOF mutations.

In conclusion, p53 R181 variants are commonly identified by germline genetic testing and confer an attenuated cancer phenotype that mimic that of other p53 hypomorphic variants. R181H and R181C maintain structural stability and tetramerization but lose the key salt-bridge with residue E180 that results in reduced DNA binding cooperativity contributing to poor binding to target promoter sites and defective transcriptional activity. As observed in other hypomorphic variants, the R181 variants retain transcription-independent function at the mitochondria to induce apoptosis. Given the levels of phenotypic heterogeneity between different p53 variants, it is critical to determine the effect of individual mutations on specific p53 tumor suppressive functions to improve our understanding of the clinical presentation and thus screening recommendations for carriers of hypomorphic TP53 variants. An understanding of specific defects of individual TP53 variants is also important for acquired mutations as *TP53* mutation status becomes increasingly important as a prognostic and predictive biomarker.

## Methods

### Patient Cohort Analysis

Analysis of clinical data and human samples was approved by the Institutional Review Boards of the University of Pennsylvania (Penn) and Children’s Hospital of Philadelphia (CHOP) (Penn IRB#834147/CHOP IRB#18-015810), Dana Farber Cancer Institute (IRB#20- 226/IRB#21-694), and Huntsman Cancer Institute (IRB#00046740). Inclusion criteria for human studies included a confirmed germline mutation in *TP53*; both males and females were included in the study. Subject demographics, cancer types, and ages at cancer diagnoses were ascertained from families with *TP53* c.542G>A;p.R181H and *TP53* c.541C>T;p.R181C across three academic medical centers (Penn, DFCI, and HCI) (**Table S1**). After exclusion of duplicate families and families with other high-risk mutations, the phenotype characteristics among p.R181H/C carriers were compared to that of carriers with other pathogenic/likely pathogenic variants in *TP53* whose data was collected from the same medical centers. Variants were categorized into hypomorphic, dominant negative/loss of function (DN/LOF) mutation classes using Kato(45) and Giacomelli(46) functional assay information in the NCI *TP53* database(33) as per ClinGen guidelines(47). The first cancer type, age at first cancer, breast cancer, and age at first breast cancer were obtained for all carriers. For those without a cancer history, age at last follow up and age at death were collected if available. Additionally, phenotype characteristics using Classic(48) and Revised Chompret(49) criteria were assessed for all variant carriers, and LFS tumor spectrum classifications(26) were assigned accordingly.

### Antibodies, cell lines, drugs, software

Information including vendor, catalog number and RRID for antibodies, cell lines, drugs, and software, where applicable, used in this study are found in **Table S12**. All cell lines were confirmed for genotype, negative for mycoplasma and with STR profiles matching those of parental lines with the Penn Genomics and Sequencing Core (RRID:SCR_024999). Single cell clones of HCT116, MCF7, and LNCaP cell lines (ATCC) with *TP53* +/-, -/-, R181H/-, and R181C/- mutations were generated with CRISPR engineering (**Table S13**) at the Washington University Genome Engineering & Stem Cell Center (RRID:SCR_023243).

### Immunoblotting

Cells were lysed using RIPA buffer and proteins quantified using BCA (Bio-Rad). 10-50μg of protein lysates were subjected to Western blotting using following antibodies overnight at 4°C: 1:1000 p53 (DO-1), 1:5000 p21 (12D1), 1:1000 MDM2 (D1V2Z), 1:750 Cleaved Lamin A (Small subunit), 1:1000 Puma, 1:10,000 GAPDH (14C10). Mouse or rabbit IgG secondary antibodies conjugated to horseradish peroxidase (Jackson ImmunoResearch) were used, followed by treatment with ECL (Millipore Sigma) for 4 minutes. Bands were imaged using fluorescence detection through the gel documentation system

### Human tumor analyses

p53 immunohistochemistry (IHC) was performed at the Penn Tumor Tissue Biospecimen Bank Core Facility (RRID:SCR_022430). Five micron sections of formalin-fixed paraffin-embedded tissue were stained using p53 antibody DO-7, diluted at 1:60. Staining was done on a Leica Bond-IIITM instrument using the Bond Polymer Refine Detection System (Leica Microsystems DS9800). Heat-induced epitope retrieval was done for 20 minutes with ER1 solution (Leica Microsystems AR9961). The presence of genomic locus-specific LOH in *TP53* was determined as previously described(28). Briefly, tumor DNA from formalin-fixed paraffin-embedded (FFPE) tumor blocks and matched germline blood DNA from LFS patients were sequenced on a NovaSeq6000 at the CHOP Center for Applied Genomics and analyzed using Varscan2 and Sequenza version 3.0.0(50). If the observed somatic variant allele fraction (VAF) exceeded the expected LOH VAF and/or the ‘B allele’ copy number was zero, the variant was classified as having locus-specific LOH.

### Protein Purification

Site-directed mutagenesis (QuikChange II; Agilent) was performed on 94-312 PET (p53 DNA-Binding Domain) plasmids (Addgene) to create p53 R181H and R181C plasmids. WT, R181H, and R181C plasmids were expressed in BL21(DE3) competent cells which were cultured overnight, lysed, and subjected to strong cation exchange chromatography. Protein was eluted and fractions containing p53 constructs were pooled and concentrated.

### Fluorescence Polarization (FP) Assay

Fluorescently labeled CDKN1A–5′-site at 1 μmol/L and unlabeled competitor sequences (**Table S13**) at 1mmol/L were reannealed at 95°C for 5 minutes and then decrease at 1°C/minute until 20°C. Apparent K_D_ values were determined by titrating WT, R181H, and R181C purified DBDs each against constant labeled CDKN1A 5′-site DNA (1nmol/L final concentration) in 50mmol/L NaPi (pH 7.2), 10% glycerol, and 1 mmol/L TCEP at 25°C for 1 hour before reading. Competition assays were conducted by premixing 2×stocks for p53/p21–5′-site DNA (28.2nmol/L for WT, 73nmol/L for R181H, and 695.6nmol/L for R181C) corresponding to 1.75x apparent K_D_ in the presence of 2nmol/L labeled CDKN1A 5′-site DNA and incubated at 25°C for 30 minutes. Competition oligonucleotides were then subjected to serial dilutions at a 2× concentration in 100 μL volume in black opaque flat bottom plates (Greiner Bio-One 655209).

Preincubated p53/CDKN1A 5′-site DNA was then combined in a 1:1 ratio with competition oligo dilutions for a final well volume of 200μL and incubated at 25°C for 1 hour. Data was collected using 458nm excitation and 528nm emission filters on a SpectraMax i3x microplate reader (Molecular Devices). All curves were fit using shared parameters for upper and lower baselines for WT, R181H, and R181C respectively (GraphPad Prism).

### Differential Scanning Fluorimetry

p53 WT, R181H, and R181C DBDs were combined at 10μmol/L each of with 10×SYPRO Orange (Invitrogen) in an assay buffer of 50 mmol/L NaPi (pH 6.3), 100 mmol/L NaCl, and 1 mmol/L DTT. Samples were heated at a rate of 1°C/second from 25°C to 65°C in an Eppendorf Realplex2 Mastercycler and monitored using the JOE emission filter (550 nm). Samples were run in four replicates. The protein’s apparent melting temperature was determined from each curve as the midpoint between the highest and lowest fluorescence values in the unfolding curve.

### Quantitative PCR (qPCR) and RNA-sequencing

HCT116 and MCF7 cells were seeded at 1-2×10^6^ cells per condition in triplicate and treated with vehicle control, 10 μmol/L Nutlin-3a (Sigma) for 0, 8, and 24 hours. RNA was isolated using QIAshredder columns (Qiagen) and the RNeasy Mini Kit (Qiagen). For qPCR, 2mg of RNA per sample (3 biological replicates per condition) was subjected to the High-Capacity Reverse Transcription Kit (Applied Biosystems). qPCR was performed using PowerUp SYBR Green Master Mix and the indicated primer sets (**Table S13**) on a QuantStudio 5 Real-Time PCR System (Applied Biosystems). Gene expression levels were normalized relative to *GAPDH* expression using the comparative Ct method. Gene expression data was expressed as relative mRNA quantity to untreated WT controls. For RNA-sequencing, libraries were prepared using the Illumina Stranded Total RNA Prep, Ligation with Ribo-Zero Plus Kit (Illumina). Overall library size was determined using high sensitivity Qubit, and Agilent Bioanalyzer (Agilent).

Libraries were pooled and subjected to paired-end next generation sequencing on the NovaSeq6000 (Illumina). FASTQ files were aligned to human genome version 19 (hg19) using STAR version 2.4.1d(51) and transcripts were mapped using Ensembl transcriptome information. Differential expression analysis of genes was performed using DESeQ2(52) for R181H/- and R181C/- cell lines against WT/-, and for differences in response to Nutlin-3a treatment. Genes less responsive to treatment in R181-mutant cells (log(fold change)<1) than in WT cells that passed a threshold of FDR<5% were used for heat maps and as inputs for enrichment analysis using IPA software (Qiagen)) using “Canonical Pathway” and “Upstream Regulator” options where a z-score below -2 were considered significantly repressed.

### Flow Cytometry

HCT116 cells were seeded at a density of 1×10^6^ cells per condition (untreated, 10 μmol/L Nutlin-3a, 10 μmol/L 5-FU in triplicate and harvested after 24 and/or 48 hours. For propidium iodide (PI) staining, cells were fixed with ethanol and processed through the Propidium Iodide Flow Cytometry Kit for Cell Cycle Analysis (abcam) according to manufacturer’s protocol. For Annexin V/PI staining, cell pellets were directly subjected to the BD Pharmingen FITC Annexin V apoptosis Detection Kit I (BD Biosciences). FITC conjugated cytochrome C release was analyzed by flow cytometry on an BD Accuri C6 Plus and analyzed using the BD Accuri software.

### Chromatin Immunoprecipitation-Sequencing (ChIP-seq)

ChIP-seq was performed in two biological replicates. For each biological replicate, 2×10^7^ of HCT116 cells were treated with vehicle control (DMSO) or 10μmol/L Nutlin-3a for 24 hours and cross-linked with 1% formaldehyde for 5 minutes at room temperature, and subsequently quenched with 0.125 M glycine for 5 minutes. Cross-linked cells were then washed twice in ice-cold 1X PBS, collected by centrifugation (2,000×*g* for 5 minutes), and lysed in 10 mL swelling buffer [10 mmol/L Tris (pH 7.5), 2 mmol/L MgCl_2_, 3 mmol/L CaCl_2_] and placed on ice for 20 minutes. Chromatin fragmentation was performed with a sonicator at 4°C with the following settings: six 10-second pulses at 20% amplitude, followed by six 10-second pulses at 25% amplitude with 10-second pauses in between pulses. Chromatin was checked for fragmentation by isolating 2% of sonicate for Qiaquick PCR purification (Qiagen, 28104) and DNA gel electrophoresis. Chromatin with sufficient shearing (200-900 base pairs) was then cleared with centrifugation at maximum speed for 10 minutes at 4°C, 5% saved for input, and immunoprecipitated with antibodies for p53 (DO-1), and RNA Polymerase II overnight at 4°C using magnetic Protein G beads (Dynabeads 10003D, Thermo). Beads were subjected to two five-minute washes with ChIP buffer [50 mmol/L HEPES (pH 7.5), 155 mmol/L NaCl, 1.1% Triton X-100, 0.11% Na-deoxycholate, 1 mmol/L EDTA], a 5-minute wash with ChIP buffer with additional 500 nmol/L NaCl, and a 5-minute wash with TE buffer [10 mmol/L Tris-HCl (pH 8.0), 1 mmol/L EDTA]. The cross-link was reversed overnight at 65°C with elution buffer [50 mmol/L Tris-HCl (pH 8.0), 10 mmol/L EDTA, 1% SDS] before treatment with 0.5 mg/mL Proteinase K (EO0491, Thermo) at 65°C for one hour. DNA was isolated and purified with the Zymo ChIP DNA Clean & Concentrator (D5205). DNA libraries were prepared using NEBNext Ultra II DNA Library Preparation kit (E7103) with NEBNext Multiplex Oligos for Illumina (7335S/L). Libraries were pooled and subjected to paired-end next generation sequencing on the NovaSeq6000 (Illumina) with 50 base pair reads. Sequences were aligned to human reference hg19 using BWA Samtools (1.9.0) (53,54) was used to remove the PCR duplicates (rmdup) and the reads with a mapping quality score of less than 10 from the aligned reads. Bigwig files of the data generated with deeptools (v2.4.2, bamCoverage–binSize 10–normalizeTo1× 3137161264– extendReads 150–ignoreForNormalization chrX(55)) and visualized on the UCSC Genome Browser. For normalization of the data, each number of the filtered reads was divided by the lowest number of the filtered reads in the same set of experiments, generating a downsampling factor for each sample. Normalized BAM files were generated using samtools view -s with the above downsampling factors and further converted to normalized BAM files using bamCoverage–binSize 10–extendReads 150. Significant peaks were called by MACS2(56) with a q-value<0.05. Heatmaps were generated from read depth normalized bigwig files using deeptools ComputeMatrix and visualized with plotHeatmap. Motif analysis was done using HOMERv4.10.1 findMotifsGenome command and known motifs were plotted.

### Mitochondrial Fractionation

HCT116 cells were seeded at 30×10^6^ cells per condition (untreated, 10 μmol/L 5-fluorouracil (5FU)) and harvested after 24 hours. Cells were washed with cold 1X PBS and subjected to fractionation buffer (0.1M Tris-MOPS, 0.1 M EGTA/Tris, 1M sucrose, and protease inhibitors). Cells were dounce homogenized using a Teflon pestle at 1,600 rpm for 60 strokes and subjected to centrifugation to isolate mitochondrial and cytosolic fractions.

### Colony Suppression Assay

Site-directed mutagenesis (QuikChange; Agilent) was performed on CMV pcDNA3.1 plasmids (Invitrogen) coding full-length TP53 to create R181H and R181C plasmids. H1299 cells were transfected with the indicated plasmid using Lipofectamine LTX and Plus Reagent (Invitrogen). After 24 hours, cells were plated at 0.5-5×10^4^ cells in triplicate. Cells were selected in 800μg/mL Geneticin (Gibco). A nontransfected control was performed. After observation of single colonies (∼7 days), cells were fixed in 10% formalin (Research Products International) for 10 minutes and stained with 0.5% crystal violet (Sigma). Plates were imaged and single colonies were counted using ImageJ.

### Mouse Experiments

All mice were treated in accordance with the regulatory standards of the NIH and American Association of Laboratory Animal Care and were approved by the Wistar Institution of Animal Care and Use Committee (IACUC; Animal Welfare Assurance ID A3432-01). Unless otherwise noted, animal experiments had equal representation of male and female mice. Food and water were provided *ad libitum*. All mice were NSG mice acquired from The Wistar Institute Animal Facility. HCT116 cells (WT/-, and two clones each of R181H/-, and R181C/-) were subcutaneously injected into the flanks of 8- to 10-week-old NSG SCID mice at a concentration of 1×10^6^ cells in 200μL 1X PBS (6 mice per cell line). Tumors were measured using a caliper 3-times a week after reaching ∼50 mm^3^ and volumes were calculated. Mice were sacrificed and tumors weighed once the tumor area reached ∼1-to 1.5-cm diameter in at least one mouse.

### Immunofluorescence (IF) and Proximity Ligation Assays (PLA)

IF and PLA experiments were performed on cells grown on Lab-Tek II 4-well or 8-well chamber slides (Thermo Fisher) and were either untreated or treated with 10μmol/L 5-Fluorouracil (5FU) or 10 μmol/L Nutlin-3a for 24 hours. Cells were fixed in 4% paraformaldehyde for 10 minutes, followed by three 5-minute washes in 1X PBS and permeabilization with 0.5% Triton X-100 (Millipore Sigma) for 10 minutes. For IF staining, cells were incubated overnight at 4°C with Ki67 and Mouse G3A1 mAb IgG1 Isotype Control diluted 1:200 in 5% goat serum (Jackson ImmunoResearch) blocking buffer. After three 5-minute 1xPBS washes, cells were incubated with Alexa Fluor 488 AffiniPure Goat anti-Mouse IgG (Jackson ImmunoResearch) diluted 1:400 in 5% goat serum blocking buffer at room temperature for 1 hour. Cells were mounted with media containing DAPI (Millipore Sigma). Ki67 images were captured using the Nikon Ti Microscope analyzing 100-250 cells per slide. p53-BAK interactions were measured using the PLA Duolink *in situ* starter kit (Sigma-Aldrich), including single-antibody controls.

Images were captured using the Leica SP8 confocal microscope. PLA was quantified by the number of signals per cell in n=10 random fields of view (>100 cells).

### Statistical Analysis

All assays were performed in biological triplicates and analyzed by unpaired two-tailed students t test (to compare between two groups such as WT vs R181H) and ANOVA (to compare between multiple groups). Kaplan–Meier survival curves were generated and analyzed using the log-rank (Mantel–Cox) test. Cancer distribution patterns between different p53 variants in carriers of different ages were statistically analyzed using the chi-squared test. All statistical analyses were performed on GraphPad Prism, with RNA-sequencing analysis performed in R. *, *P* < 0.05; **, *P* < 0.01; ***, *P* < 0.001; **** *P* < 0.0001; ns, not significant.

## Acknowledgements

Research support for this study was provided by The National Institutes of Health (F31CA277953, AI; U54CA272686, SAM, JK; P30CA042014, LDM; R01CA242218, JEG; R01CA102184, MEM; K08CA215312, KNM); the Li Fraumeni Syndrome Association (KNM); the Burroughs Wellcome Fund (1017184, KNM); and the Basser Center for *BRCA* at the University of Pennsylvania (RH, KNM). We thank the Genome Engineering & Stem Cell Center (GESC@MGI) at Washington University in St. Louis for cell line engineering services. We thank the Wistar Microscopy Core for their expertise in immunofluorescence and proximity ligation assay image capture and analysis. We thank the Cancer Research Shared Resource and Hyundai Hope on Wheels for HCI’s contribution.

## Data Availability Statement

It is not possible for the authors to directly share the individual-level clinical data that were obtained due to IRB constraints. Anyone wishing to gain access to this data should inquire directly with the authors. The data generated from our analyses are included in the manuscript main text, tables, and figures and online Supplementary Materials (available online). The sequencing data is deposited in the Gene Expression Omnibus (GEO) repository under accession number GSE303326. Tumor genomic sequencing data is deposited in the dbGaP database entitled LFS Genomics under accession number phs003348.

## Authorship Contributions

Conception/design: RM, JK, MEM, KNM; Provision of study material or patients: ASL, IA, MH, RD, CO, WK, AN, JV, SRC, LDM, JEG, KNM; Collection and/or assembly of data: RM, ASL, GK, SAM, IA, MH, RD, CO, WK, AN, JV, SRC; Data analysis and interpretation: RM, AI, RH, ASL, SAM, MEM, KNM; Manuscript writing: RM, AI, RH, ASL, MEM, KNM; Final approval of manuscript: All authors.

