## Supplemental Data for "Variation at the R181 residue of p53 confers loss of p53 DNA binding cooperativity with the retention of mitochondrial-associated apoptosis"

**Supplement Contents**

1. **Supplemental Table 1:** Detailed clinical information on R181H/C probands from three academic medical centers
2. **Supplemental Table 2:** History of cancer and age of onset in carriers of R181, DN/LOF, and hypomorphic variants in the patient cohort
3. **Supplemental Table 3:** Kratz criteria assigned to carriers of R181, DN/LOF, and hypomorphic variants in the patient cohort
4. **Supplemental Table 4:** Differential expression analysis of 328 p53 target genes in HCT116 p53 +/-, R181H/-. R181C/- cells after 8 hours of Nutlin-3a compared to vehicle
5. **Supplemental Table 5:** Differential expression analysis of 328 p53 target genes in HCT116 p53 +/-, R181H/-. R181C/- cells after 24 hours of Nutlin-3a compared to vehicle
6. **Supplemental Table 6:** Ingenuity Pathway Analysis of RNAseq data after 8hr of Nutlin-3a in HCT116 p53 R181H/- vs +/- cells
7. **Supplemental Table 7:** Ingenuity Pathway Analysis of RNAseq data after 8hr of Nutlin-3a in HCT116 p53 R181C/- vs +/- cells
8. **Supplemental Table 8:** Differential expression analysis of 328 p53 target genes in MCF7 p53 +/-, R181H/-. R181C/- cells after 8 hours of Nutlin-3a compared to vehicle
9. **Supplemental Table 9:** Differential expression analysis of 328 p53 target genes in MCF7 p53 +/-, R181H/-. R181C/- cells after 24 hours of Nutlin-3a compared to vehicle
10. **Supplemental Table 10:** Ingenuity Pathway Analysis of RNAseq data after 8hr of Nutlin-3a in MCF7 p53 R181H/- vs +/- cells
11. **Supplemental Table 11:** Ingenuity Pathway Analysis of RNAseq data after 8hr of Nutlin-3a in MCF7 p53 R181C/- vs +/- cells
12. **Supplemental Table 12**
    a. **12A:** Antibodies, cell lines, and drugs used in this study
    b. **12B:** Software and algorithms used in this study
13. **Supplemental Table 13:** Oligonucleotides used in this study
14. **Supplemental Figure 1.** *TP53* c.542G>A;p.R181H and *TP53* c.541C>T;p.R181C variants are the most common variants observed at Penn, DFCI, and HCI and confer the risk of various tumors.
15. **Supplemental Figure 2.** R181H Pedigrees from Penn Medicine (Penn), Dana-Farber Cancer Institute (DFCI), and Huntsman Cancer Institute (HCI).
16. **Supplemental Figure 3.** R181H tumors undergo loss of heterozygosity.
17. **Supplemental Figure 4.** RNA-sequencing and quantitative PCR of R181-mutant cell lines confirm reduced induction of p53 target genes.
18. **Supplemental Figure 5.** Second replicate of ChIP-sequencing and HOMER motif analysis showing retained binding of R181-mutants to ETS sites.
19. **Supplemental Figure 6.** R181-mutant HCT116 tumors grow similar to WT tumors in the flanks of mice.
20. **Supplemental Figure 7.** R181 variants do not have aberrant mitochondrial function.

**Supplementary Tables**

*See attached Excel File*

**Supplementary Figure Legends**

**Supplemental Figure 1.** *TP53* c.542G>A;p.R181H and *TP53* c.541C>T;p.R181C variants are the most common variants observed at Penn, DFCI, and HCI and confer the risk of various tumors. (A) Pie chart showing proportion of germline *TP53* variants observed in families (n= 145) at Penn Medicine (Penn). (B) Pie chart showing proportion of germline *TP53* variants observed in families (n= 174) at Dana-Farber Cancer Institute (DFCI). (C) Pie chart showing proportion of germline *TP53* variants observed in families (n= 96) at Huntsman Cancer Institute (HCI). (D) Consort diagram describing the R181H and R181C cohort studied for clinical analysis. (E) Bar chart showing total tumor distribution (primary and other malignancies) in R181-variant carriers from Penn, DFCI, and HCI (n= 109 tumors). (F) Bar plot showing R181-mutant tumor types identified in AACR GENIE consortium which does not differentiate between germline and somatic R181 mutations. (G) Bar plot showing the percentage of R181-carriers with different primary cancer diagnoses. Breast, cervical, ovarian, and endometrial cancer reported in a separate bar plot as a percentage of female patients. Prostate and testicular cancer reported in a separate bar plot as a percentage of male patients. (H) Bar plot showing the percentage of Hypomorphic variant carriers with different primary cancer diagnoses. Breast, cervical, and endometrial cancer reported in a separate bar plot as a percentage of female patients. Prostate and testicular cancer reported in a separate bar plot as a percentage of male patients. Variants included as hypomorphic were: p.G105D, p.R110C, p.R110L, p.T125M, p.N131K, p.G154D, p.R156H, p.R158C, p.A161T, p.R175C, p.H178Q, p.P190R, p.H193Q, p.H214Y, p.V218M, p.R267Q, p.R267W, p.C277R, p.D281E, p.R282Q, p.G245C, p.G334R, p.A347D. (I) Bar plot showing the percentage of DN/LOF carriers with different primary cancer diagnoses. Breast, cervical, and ovarian cancer reported in a separate bar plot as a percentage of female patients. Prostate cancer reported in a separate bar plot as a percentage of male patients.

**Supplemental Figure 2.** Representative Pedigrees of families with the *TP53* p.R181H pathogenic germline variant from Penn Medicine (Penn), Dana-Farber Cancer Institute (DFCI), and Huntsman Cancer Institute (HCI). Probands are indicated by the black arrowhead; slashes indicated deceased status. Circles represent females, squares represent males. Numbers under symbols indicate current age or age at death. Cancer diagnoses are indicated by a black quarter fill with abbreviation for cancer diagnosis and age at diagnosis under the individual. ACC, adrenocortical cancer; BR, breast cancer; CNS, central nervous system cancer; CO, colon cancer; CX, cervical cancer; DCIS, ductal carcinoma in situ; HN, head and neck cancer; LG, lung cancer; LK, Leukemia; MEL, melanoma; URT, ureter cancer, MM, multiple myeloma; PAN, pancreatic cancer; PR, prostate cancer; PTC, papillary thyroid cancer; STO, stomach cancer; THY, thyroid cancer.

**Supplemental Figure 3.** R181H tumors undergo loss of heterozygosity. Genome view of copy number alterations data generated by applying the sequenza package to sequencing of tumor DNA from FFPE tumor blocks from patients with germline *TP53* variants (A) p.G245S, (B) p.R248Q, and (C) – (D) p.R181H shows loss of heterozygosity at the *TP53* locus in chromosome 17 (17p13.1). DCIS, Ductal Carcinoma in situ; IDC, Invasive Ductal Carcinoma.

**Supplemental Figure 4.** RNA-sequencing and quantitative PCR of R181-mutant cell lines confirm reduced induction of p53 target genes. (A) Volcano plots of differentially expressed genes in HCT116 R181H/- cells compared to p53 WT/- cells and HCT116 R181C/- cells compared to p53 WT/- cells after 8 hours of Nutlin-3a treatment. (B) Heatmap showing log2(Fold Change) after 8 hours of vehicle or Nutlin-3a in HCT116 p53 WT/- cells and R181H/- cells; genes shown are p53 targets that were significantly up-regulated in WT/- cells (n=48). Bar to the right of the heatmap depicts the false discovery rate comparing the fold change between treatment and vehicle in HCT116 p53 WT/- versus R181H/- cells. Heatmap showing log2(Fold Change) after 8 hours of vehicle or Nutlin-3a in HCT116 p53 WT/- cells and R181C/- cells; genes shown are p53 targets that were significantly up-regulated in WT/- cells (n=48). Bar to the right of the heatmap depicts the false discovery rate comparing the fold change between treatment and vehicle in HCT116 p53 WT/- versus R181H cells. (C) Bar plot showing the -log(p-value) of the most significantly decreased regulators in HCT116 R181H/- vs WT/- 24 hours after Nutlin-3a treatment. Bar plot showing the -log(p-value) of the most significantly decreased regulators in HCT116 R181C/- vs WT/- 24 hours after Nutlin-3a treatment. (D) Bar plot showing delta delta Ct from quantitative PCR (qPCR) of p53 target gene *CDKN1A* in HCT116 WT/-, -/-, R181H/-, R181C/- cells. (E) Bar plot showing delta delta Ct qPCR of p53 target gene *BBC3* in HCT116 WT/-, -/-, R181H/-, R181C/- cells. (F) Bar plot showing delta delta Ct qPCR of p53 target gene *GDF15* in HCT116 WT/-, -/-, R181H/-, R181C/- cells. Data averaged from three independent experiments. Student’s t-test * P < 0.05, ** < 0.01. (G) Volcano plots of differentially expressed genes in MCF7 R181H/- cells compared to p53 WT/- cells and MCF7 R181C/- cells compared to p53 WT/- cells after 8 hours of Nutlin-3a treatment. (H) Heatmap showing log2(Fold Change) after 8 hours of vehicle or Nutlin-3a in MCF7 p53 WT/- cells and R181H/- cells; genes shown are p53 targets that were significantly up-regulated in WT/- cells (n=160). Bar to the right of the heatmap depicts the false discovery rate comparing the fold change between treatment and vehicle in MCF7 p53 WT/- versus R181H/- cells. Heatmap showing log2(Fold Change) after 8 hours of vehicle or Nutlin-3a in MCF7 p53 WT/- cells and R181C/- cells; genes shown are p53 targets that were significantly up-regulated in WT/- cells (n=160). Bar to the right of the heatmap depicts the false discovery rate comparing the fold change between treatment and vehicle in MCF7 p53 WT/- versus R181H cells. (I) Bar plot showing the -log(p-value) of the most significantly decreased regulators in MCF7 R181H/- vs WT/- 24 hours after Nutlin-3a treatment. Bar plot showing the -log(p-value) of the most significantly decreased regulators in MCF7 R181C/- vs WT/- 24 hours after Nutlin-3a treatment. (J) Western blot of p53, Mdm2, p21, and GAPDH (loading control) in WT/WT and WT/R181H lymphoblastoid cell lines (LCLs) after 0, 8, and 24 hours of Nutlin-3a treatment.

**Supplemental Figure 5.** Second replicate of ChIP-sequencing and HOMER motif analysis showing retained binding of R181-mutants to ETS sites. (A) Heatmap showing significant peaks indicating sites on DNA bound by p53 in HCT116 p53 WT/-, R181H/-, and R181C/- cells in 0, 8, and 24 hours of Nutlin-3a treatment after chromatin immunoprecipitation sequencing (ChIP-seq) (replicate 2). (B) Bar plot showing -log(p-value) of the significantly enriched motifs by p53 in HCT116 WT/-, R181H/- and R181C/- cells. Data are representative of n = 2 independent experiments.

**Supplemental Figure 6.** R181-mutant HCT116 tumors grow similar to WT tumors in the flanks of mice. (A) Growth curves of HCT116 WT/-, R181H/- clone 1, R181H/- clone 2, R181C/- clone 1, and R181C/- clone 2 cells subcutaneously injected into NSG SCID mice (6 mice per cell line). (B) Scatter plot showing final tumor weights of HCT116 WT/-, R181H/- clone 1, R181H/- clone 2, R181C/- clone 1, and R181C/- clone 2 xenografts.

**Supplemental Figure 7.** R181 variants do not have aberrant mitochondrial function. (A) Representative microscopy images of p53 and mitochondria immunofluorescence (IF) in HCT116 WT/-, R181H/-, and R181C/- cells at 0 and 24 hours of 5-fluorouracil (5FU) treatment; blue = DAPI, green = A488 (p53), red = MitoTracker (mitochondria). (B) Scatter plot of the mean fluorescence intensity (MFI) of MitoTracker measured through flow cytometry in HCT116 WT/-, -/-, R181H/-, and R181C/- cells. Data averaged from three independent experiments. (C) Scatter plot of MFI of TOMM20 measured through flow cytometry in HCT116 WT/-, -/-, R181H/-, and R181C/- cells. Data averaged from three independent experiments. Student’s t-test * P < 0.05, ** < 0.01 (D) Graph showing oxygen consumption rate (OCR) measured through Seahorse XF Mito Stress Test in HCT116 WT/-, -/-, R181H/-, and R181C/- cells. Assay performed using 12 technical replicates per cell line.

**Supplemental Figure 1**

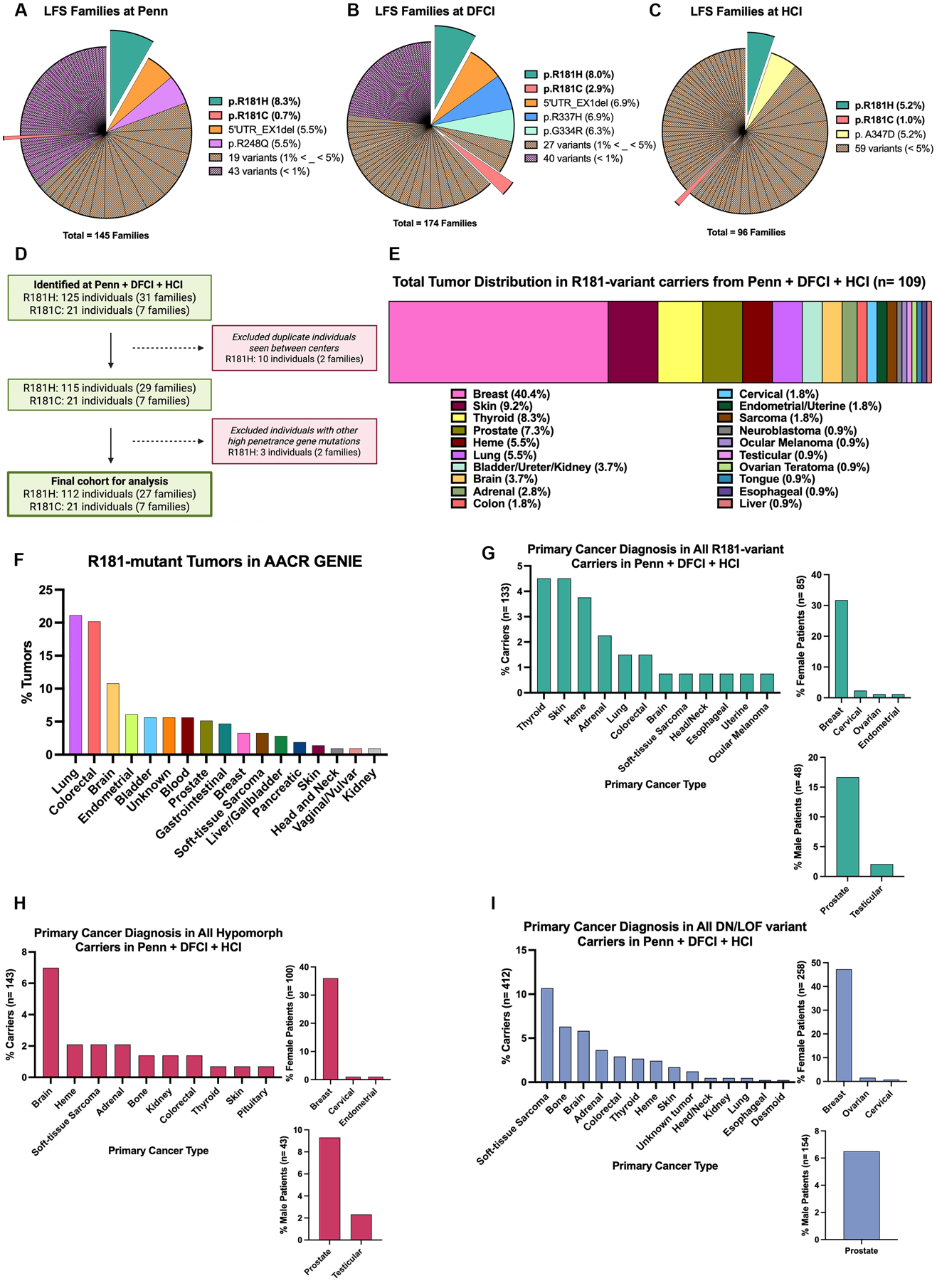

**Supplemental Figure 2**

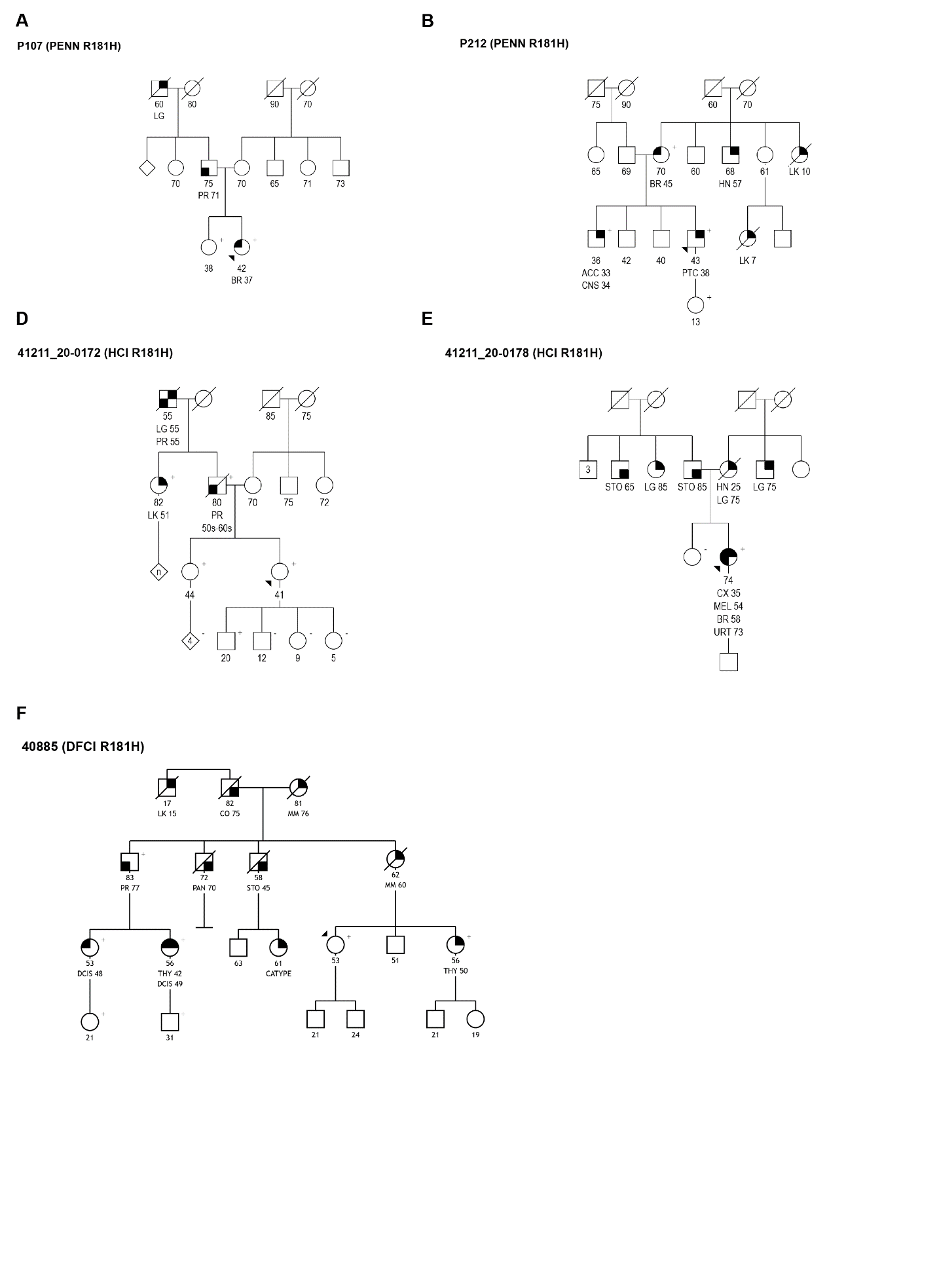

**Supplemental Figure 3**

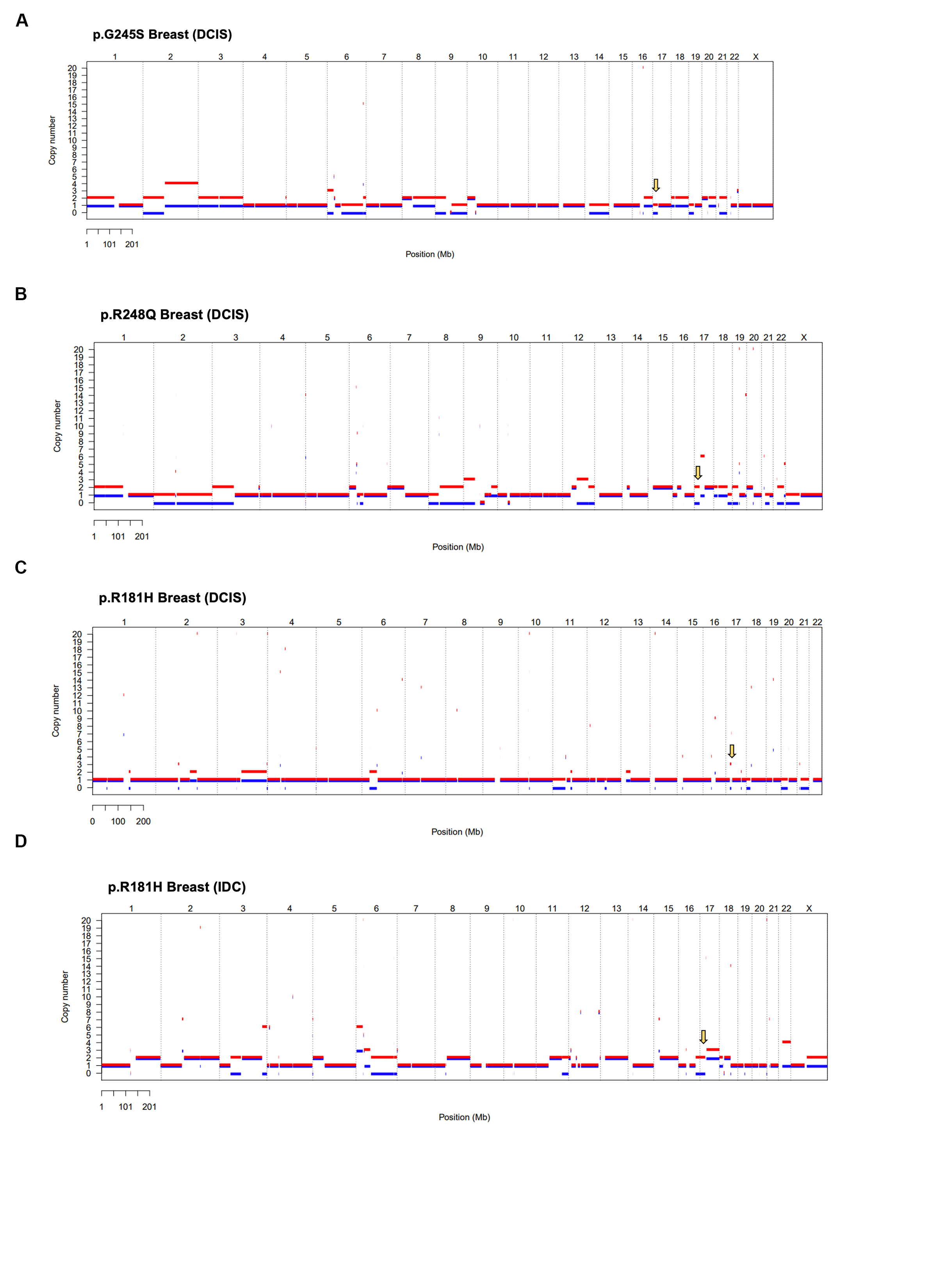

**Supplemental Figure 4**

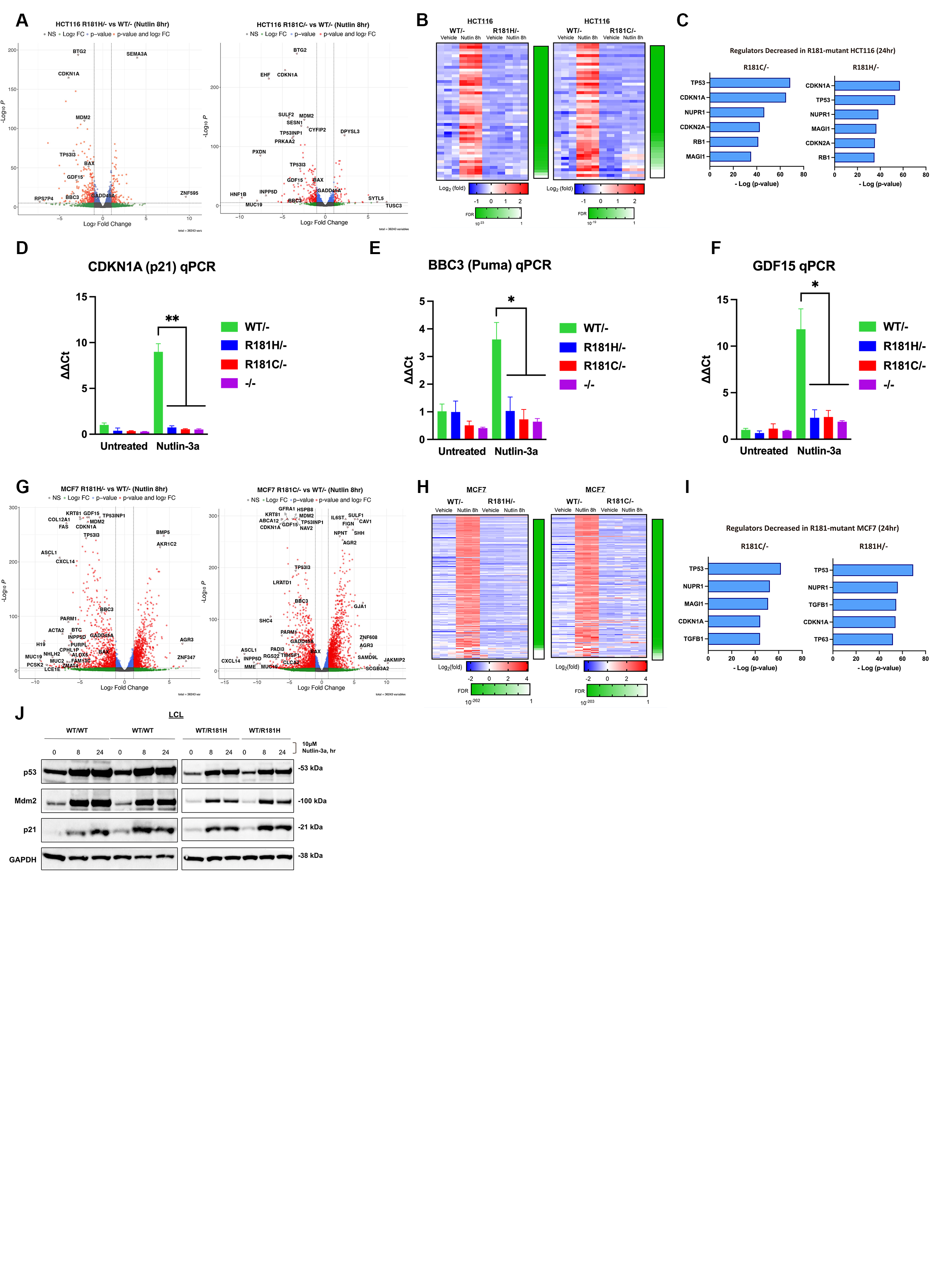

**Supplemental Figure 5**

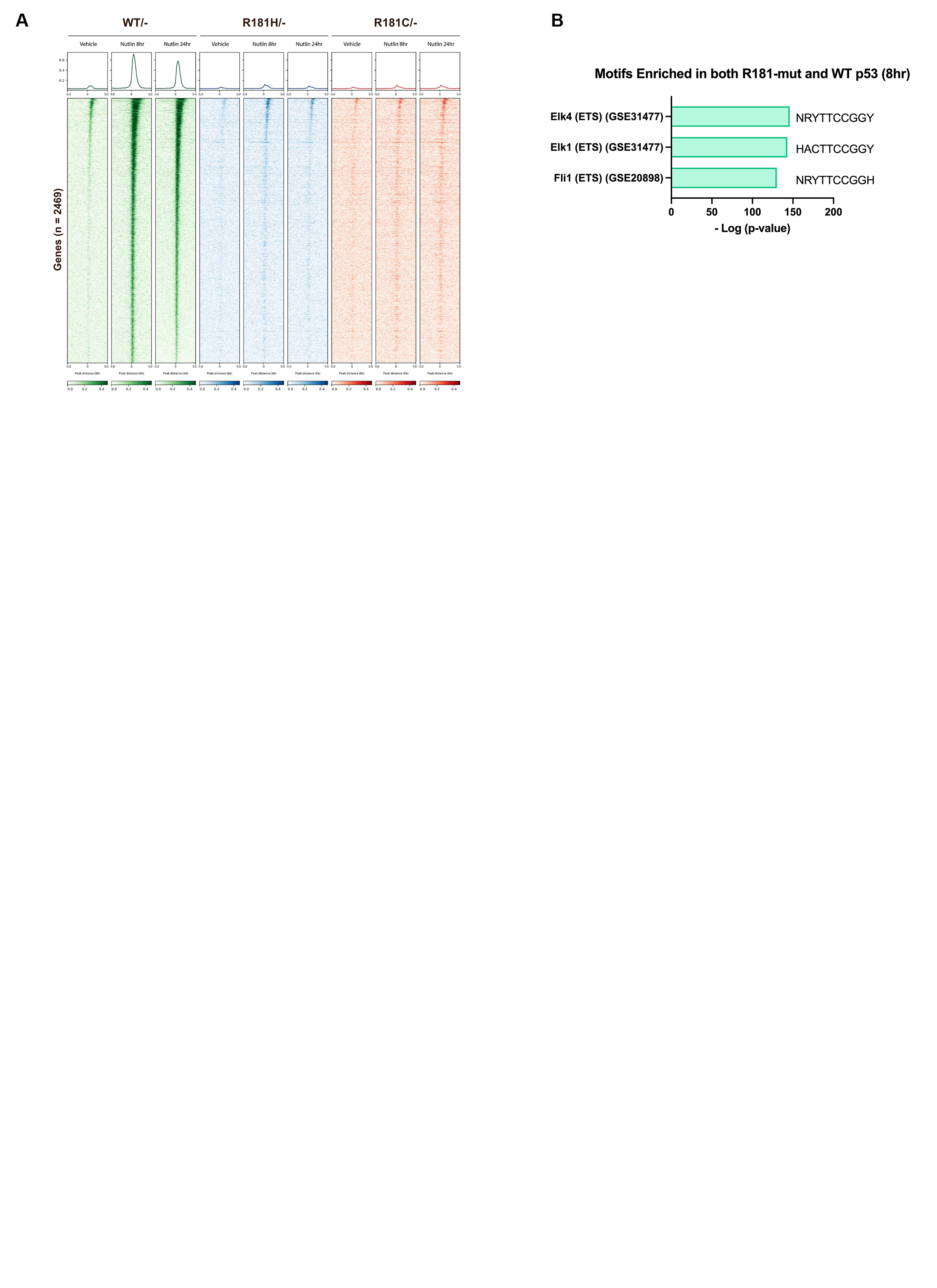

**Supplemental Figure 6**

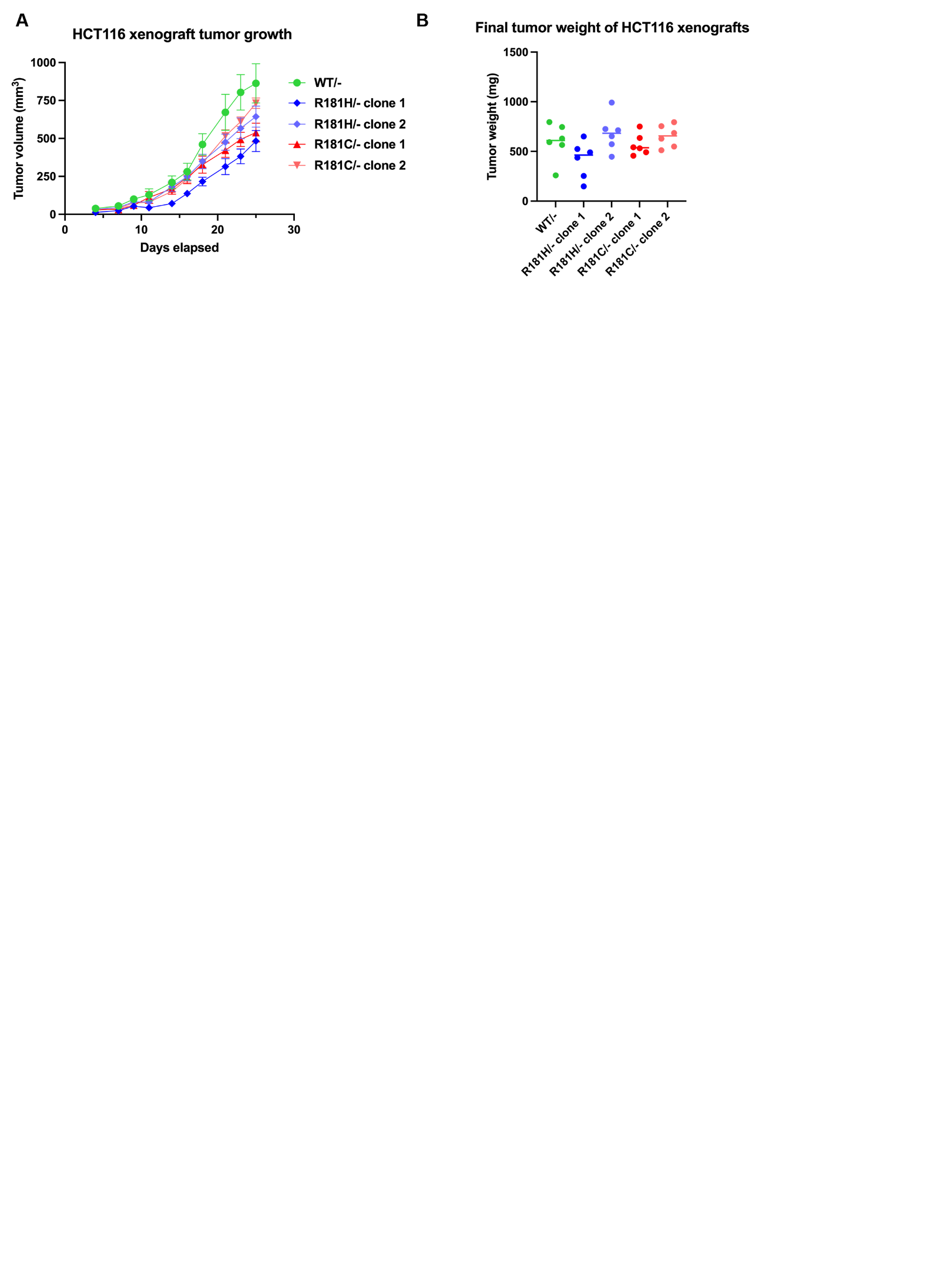

**Supplemental Figure 7**

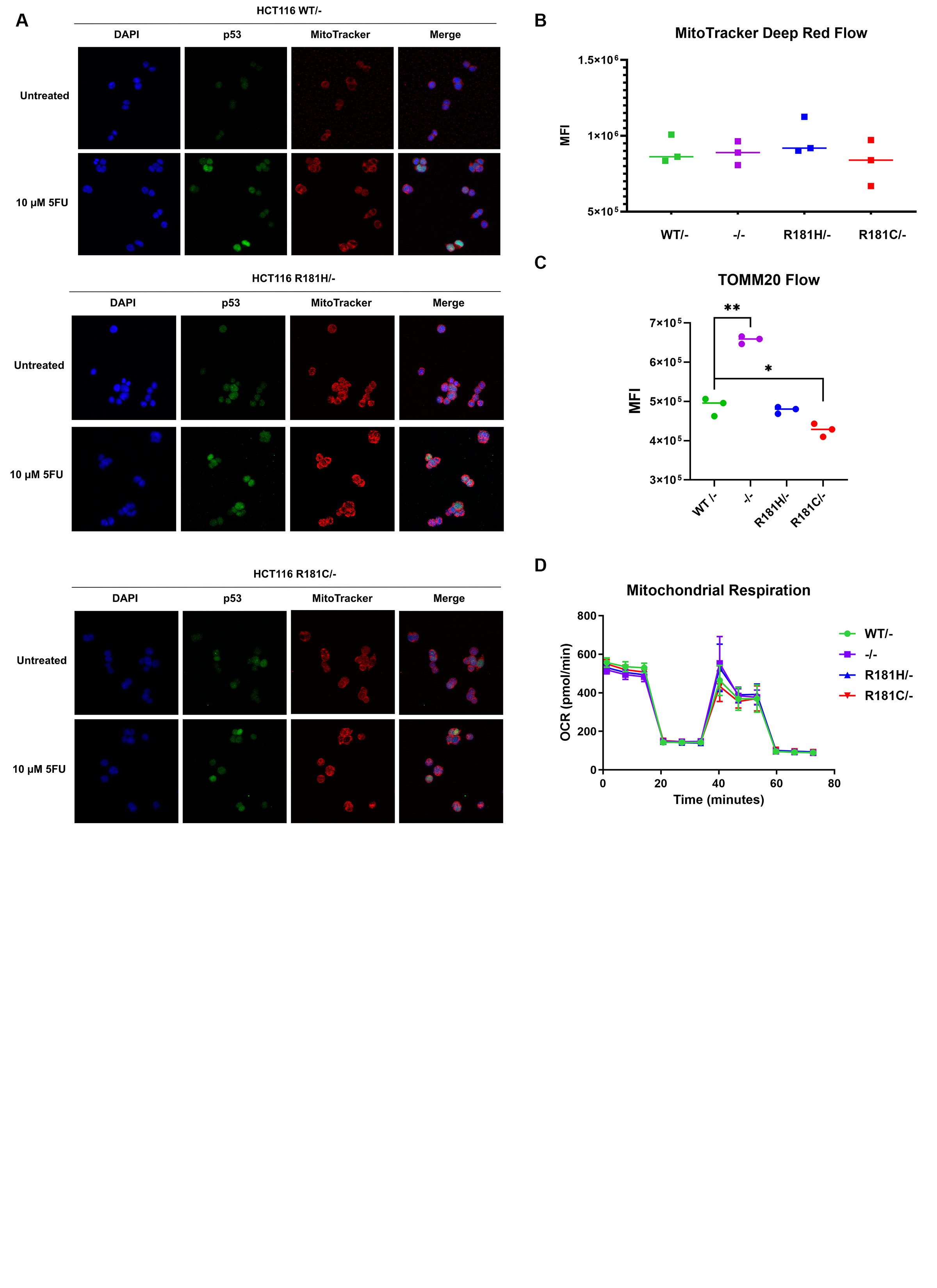
